# Lassa virus glycoprotein nanoparticles elicit a neutralizing antibody that defines a new site of vulnerability

**DOI:** 10.1101/2022.03.28.486091

**Authors:** Philip J.M. Brouwer, Aleksandar Antanasijevic, Adam J. Ronk, Helena Müller-Kräuter, Yasunori Watanabe, Mathieu Claireaux, Nicole M. Lloyd, Tom. P. L. Bijl, Hailee R. Perrett, Thijs Steijaert, Judith A. Burger, Marlies M. van Haaren, Kwinten Sliepen, Marit J. van Gils, Max Crispin, Thomas Strecker, Alexander Bukreyev, Andrew B. Ward, Rogier W. Sanders

## Abstract

Lassa virus is endemic in large parts of West Africa and causes a hemorrhagic fever. Recent years have seen several serious outbreaks of Lassa fever with high mortality rates. A vaccine to curtail infection is urgently needed. The development of a recombinant protein vaccine has been hampered by the instability of soluble Lassa virus glycoprotein complex (GPC) trimers, which disassemble into monomeric subunits after expression. Here we use two-component protein nanoparticles to stabilize GPC in a trimeric conformation and present twenty prefusion GPC trimers on the surface of an icosahedral nanoparticle. Cryo-EM studies of assembled GPC nanoparticles demonstrated a well-ordered structure and yielded a high-resolution structure of an unliganded GPC. These nanoparticles induced potent humoral immune responses in rabbits and protective immunity against a lethal Lassa virus challenge in guinea pigs. We isolated a neutralizing antibody which was mapped to the putative receptor-binding site, revealing a novel site of vulnerability on GPC.

## Introduction

Lassa Virus (LASV) is an Old World mammarenavirus and the causative agent of Lassa fever, a viral hemorrhagic fever that is endemic in parts of West Africa. The virus is shed in the urine and feces of its natural reservoir, the multimammate rat (*Mastomys natalensis)*, and spreads to humans by contact or ingestion of the rat’s excrement or saliva (Günther and Lenz, 2004). While the majority of infections are zoonotic, human-to-human transmissions, primarily in the form of nosocomial transmission, have also been reported (Andersen et al., 2015; Dan-Nwafor et al., 2019; Fisher-Hoch et al., 1995). An estimated 100,000-300,000 people are infected with LASV each year, with 5000-10,000 succumbing to the disease (McCormick et al., 1987). These numbers are however likely an underestimation due to lack of proper diagnostics in the impoverished, mostly rural, endemic areas and the nonspecific febrile symptoms of Lassa fever. Although the majority of infections are benign, for hospitalized patients the case-fatality rate is around 25% (Kenmoe et al., 2020). Recently, Nigeria has experienced several serious LASV outbreaks with around 30% of diagnosed patients not surviving the infection (Bagcchi, 2020; Kenmoe et al., 2020). Seven distinct phylogenetic lineages of LASV have been identified which cluster based on their geographic location. Lineage II and III are the most common lineages in Nigeria, whereas lineage IV is the most dominant lineage in Sierra Leone and Guinea (Ehichioya et al., 2019; Ibukun, 2020). A vaccine against LASV would ideally be able to confer protection against all currently known lineages.

Recent animal studies demonstrated that administration of neutralizing antibodies (NAbs) isolated from humans previously infected with LASV conferred 100% protection against a lethal LASV challenge, providing impetus for the development of a vaccine that induces potent NAb responses (Cross et al., 2016, 2019; Mire et al., 2017). The glycoprotein complex (GPC) is expressed as a trimer on the viral surface and constitutes the sole target for neutralizing antibodies (NAbs). Each GPC protomer consists of two non-covalently bound subunits: the membrane-anchored GP2, which contains the fusion peptide, and GP1, which possesses receptor binding sites for both alpha-dystroglycan and lysosome-associated membrane protein-1 (LAMP-1), the two receptors involved in viral entry and lysosomal escape, respectively (Hastie et al., 2017). Following the observations that stabilized prefusion glycoprotein trimers of SARS-CoV-2, HIV-1 and RSV induced strong NAb responses (Guebre-Xabier et al., 2020; McLellan et al., 2013; Pauthner et al., 2019), recombinantly expressed GPC that stably maintains a trimeric prefusion state may represent a promising immunogen to induce potent humoral immune responses against LASV. Indeed, the majority of known NAbs target prefusion GPC, of which the most potent ones target an epitope that spans multiple protomers (Hastie et al., 2019; Robinson et al., 2016). Although the development of a prefusion-stabilized GPC constituted an important first step for recombinant protein vaccine design, several challenges remain (Hastie et al., 2017). First of all, prefusion GPC trimers rapidly dissociate into monomers upon expression, resulting in the loss of NAb epitopes and exposure of immunodominant non-NAb epitopes on the non-glycosylated GPC interior (Hastie and Saphire, 2018). Second, GPC is covered by a dense glycan shield making it a poorly immunogenic antigen (Sommerstein et al., 2015; Watanabe et al., 2018). Finally, GPC is sequence diverse, necessitating the induction of bNAb responses to neutralize the several lineages of LASV, which further complicates the development of an effective pan-LASV vaccine (Ibukun, 2020).

Presenting prefusion-stabilized glycoproteins on computationally designed two-component protein nanoparticles has greatly enhanced vaccine-induced antibody responses to RSV, influenza, and HIV-1 (Boyoglu-Barnum et al. 2021; Brouwer et al., 2021a; Marcandalli et al., 2019). The most widely used two-component nanoparticle design is I53-50, which is made up of twenty trimeric (I53-50A or variants thereof) and twelve pentameric (I53-50B) subunits that self-assemble *in vitro* to form monodisperse icosahedral particles with a diameter of approximately 30 nm (Bale et al., 2016). Computational redesign of the trimeric subunit has generated an exterior-facing N-terminal helical extension on I53-50A that allows facile genetic fusion of trimeric antigens. This has enabled the generation of I53-50 nanoparticle (I53-50NP) vaccines featuring RSV-F, HIV-1 Env, and SARS-CoV-2 S glycoproteins (Brouwer et al., 2021a, 2021b; Marcandalli et al., 2019). Here, we describe the design and production of I53-50NPs that present native-like GPC trimer antigens of LASV. We applied electron microscopy (EM), biolayer interferometry (BLI) and differential scanning fluorimetry (nanoDSF) to confirm the appropriate structural and antigenic properties of the designed antigens. Finally, we assessed the immunogenicity of native-like GPC as free trimers and in the context of the nanoparticle. The GPC nanoparticles consistently induced strong antibody responses in immunized rabbits. Furthermore, vaccination with these nanoparticles protected guinea pigs from LASV-induced mortality. Finally, through analysis of memory B cells from immunized rabbits we identified a glycan-dependent NAb targeting the putative LAMP-1 binding site on GPC.

## Results

### Fusion to I53-50A improves LASV GPC trimerization

Previous work with I53-50NPs presenting HIV-1 Env and RSV-F suggested that fusion to I53-50A can have a stabilizing effect on the displayed glycoprotein (Brouwer et al., 2019; Marcandalli et al., 2019). Therefore, in an attempt to improve trimerization, we genetically fused the previously described prefusion-stabilized GPCysR4 (Josiah strain, lineage IV (Hastie et al., 2017)) to I53-50A (Fig 1A). The resulting construct, designated GPC-I53-50A for simplicity, was expressed in HEK293 cells and purified by streptactin-based affinity purification followed by size-exclusion chromatography (SEC) to yield a reasonable amount (∼ 0.7 mg/L of cells) of efficiently cleaved fusion proteins (Fig S1A). Non-scaffolded GPC was expressed poorly (< 0.1 mg/L of cells). Whereas blue-native polyacrylamide gel electrophoresis (BN-PAGE) showed no clear trimer band for non-scaffolded GPC protein, GPC-I53-50A was exclusively trimeric (Fig. 1B). 2D class-averages from negative-stain EM (nsEM) analyses revealed that the large majority of non-scaffolded GPC proteins was monomeric, while the GPC-I53-50A fusion protein was predominantly trimeric with the GPC and I53-50A components readily discernible (Fig. 1C). Fusion to I53-50A increased the thermostability of GPC from 62.7°C to 64.5°C (Fig. S1B).

**Fig. 1.**
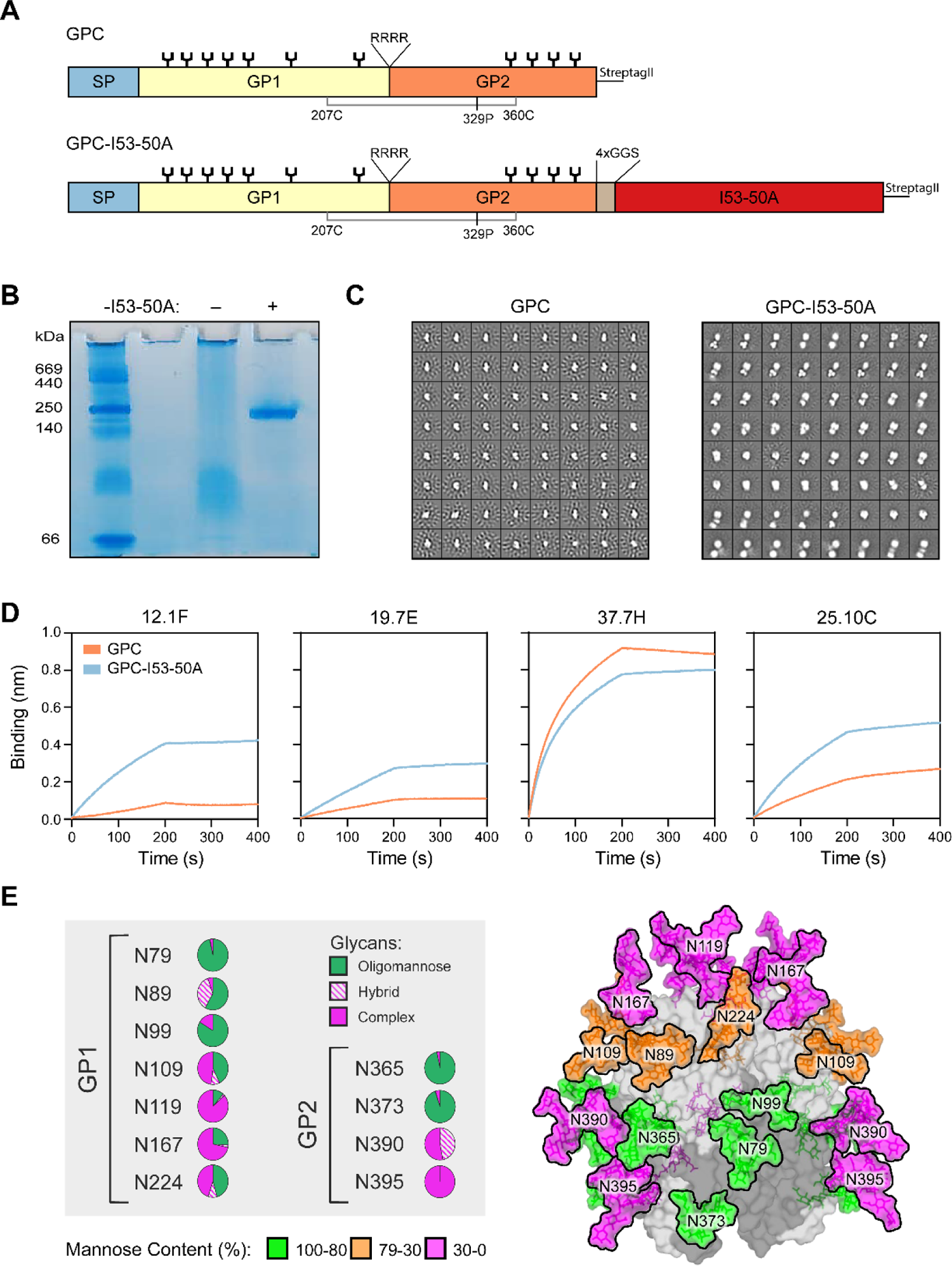
Biophysical characterization, antigenicity and glycosylation of GPC-I53-50A. (A) Linear schematic of the GPC and GPC-I53-50A constructs with GPCysR4 mutations annotated. The disulfide bond (207C-360C) that connects GP1 to GP2 is shown in grey. SP = signal peptide. (B) BN-PAGE analysis of GPC (-) and GPC-I53-50A (+). (C) 2D-class averages from nsEM with GPC (left) and GPC-I53-50A (right). (D) Sensorgrams from BLI experiments showing binding of GPC and GPC-I53-50A to immobilized human (b)NAbs 12.1F, 19.7E, 37.7H, 25.10C. (E) Site-specific glycan analysis of GPC-I53-50A. Each pie chart summarizes the quantification of oligomannose (green), hybrid (dashed pink), and complex glycans (pink) for each glycan site on GP1 or GP2. The experimentally observed glycans are modeled on the trimeric GPC structure (PDB ID: 5VK2) (Hastie et al., 2017). The glycans are coloured according to the oligomannose content as defined in the legend. GP1 and GP2 subunits are coloured light grey and dark grey, respectively.

To probe the antigenicity of GPC-I53-50A we performed bio-layer interferometry (BLI)-based binding experiments. A panel of NAbs was immobilized and binding to an equimolar amount of I53-50A-scaffolded and non-scaffolded GPC was assessed (Fig. 1D). We observed markedly improved binding to GP1-specific bNAbs 12.1F and 19.7E when GPC was fused to I53-50A. This was also the case for the potent and quaternary-dependent NAb 25.10C. Association of the NAb 37.7H, which targets an epitope that spans two protomers but also binds to monomeric GPC, was slightly lower for GPC-I53-50A than non-scaffolded GPC. However, no dissociation was observed for GPC-I53-50A, which is consistent with improved trimerization for this construct.

### LASV GPC-I53-50A presents native-like glycans

The GPC from LASV strain Josiah used to generate GPC-I53-50A encodes 11 potential N-linked glycosylation sites (PNGS) (Eichler et al., 2006). The host-derived glycans hinder antibody recognition of GPC rendering them immunologically challenging targets (Sommerstein et al., 2015; Watanabe et al., 2018). Previous analysis of the glycan shield of GPC in the context of a native-like virus-like particle (VLP) system revealed the presence of a dense cluster of oligomannose-type glycans spanning the GP1 and GP2 subunits (Watanabe et al., 2018). Since glycan modifications are sensitive reporters of protein architecture, we sought to determine the glycosylation profile of GPC-I53-50A. Glycan analysis using liquid chromatography-mass spectrometry (LC-MS) revealed a remarkable conservation in the glycan compositions between the previously described GPC on VLPs and GPC-I53-50A. Specifically, the oligomannose cluster consisting of the glycans at N79, N89, N99, N365 and N373, was conserved between the two GPC forms (Fig. 1E and fig. S1C). This underprocessing of glycans that arises due to the structural clustering of the glycans likely reflects the native-like protein architecture of GPC-I53-50A. In addition, with the exception of the glycan at N224, the remaining glycans sites were predominantly of the complex-type in both GPC-VLPs and GPC-I53-50A.

### LASV GPC-I53-50A and I53-50B efficiently assemble into icosahedral nanoparticles

To assemble icosahedral I53-50NPs presenting twenty GPC trimers, SEC-purified GPC-I53-50A was mixed with I53-50B at an equimolar ratio of monomeric subunits and incubated overnight at 4°C. The assembled nanoparticles were then separated from unassembled components by an additional SEC step. Sodium dodecyl sulfate polyacrylamide gel electrophoresis (SDS-PAGE) of the collected higher molecular weight complexes revealed that all the expected nanoparticle components were present (fig. S1A). Only minimal amounts of unassembled components were observed in SEC, suggesting efficient assembly of GPC-I53-50NPs (Fig. 2A). nsEM experiments with the pooled fractions containing high molecular weight proteins revealed a homogeneous preparation of well-ordered icosahedral particles, with a small percentage of aggregated GPC-I53-50NPs. The latter was confirmed by dynamic light scattering (DLS), which showed a monomodal but slightly polydisperse population of particles (Fig. 2B and fig. S1D). Similar to our observations with I53-50NPs presenting native-like HIV-1 Env trimers (Brouwer et al., 2019, 2021a), NanoDSF showed a two-phased melting curve for nanoparticles, with a melting temperature for GPC at 63.6°C and for the I53-50 components at 81°C. The GPC-I53-50NPs retained the capacity to bind to the monoclonal bNAbs 19.7E, 12.1F, 37.7H and 25.10C confirming that nanoparticle assembly did not compromise the antigenicity of GPC (Fig. 2C).

**Fig. 2.**
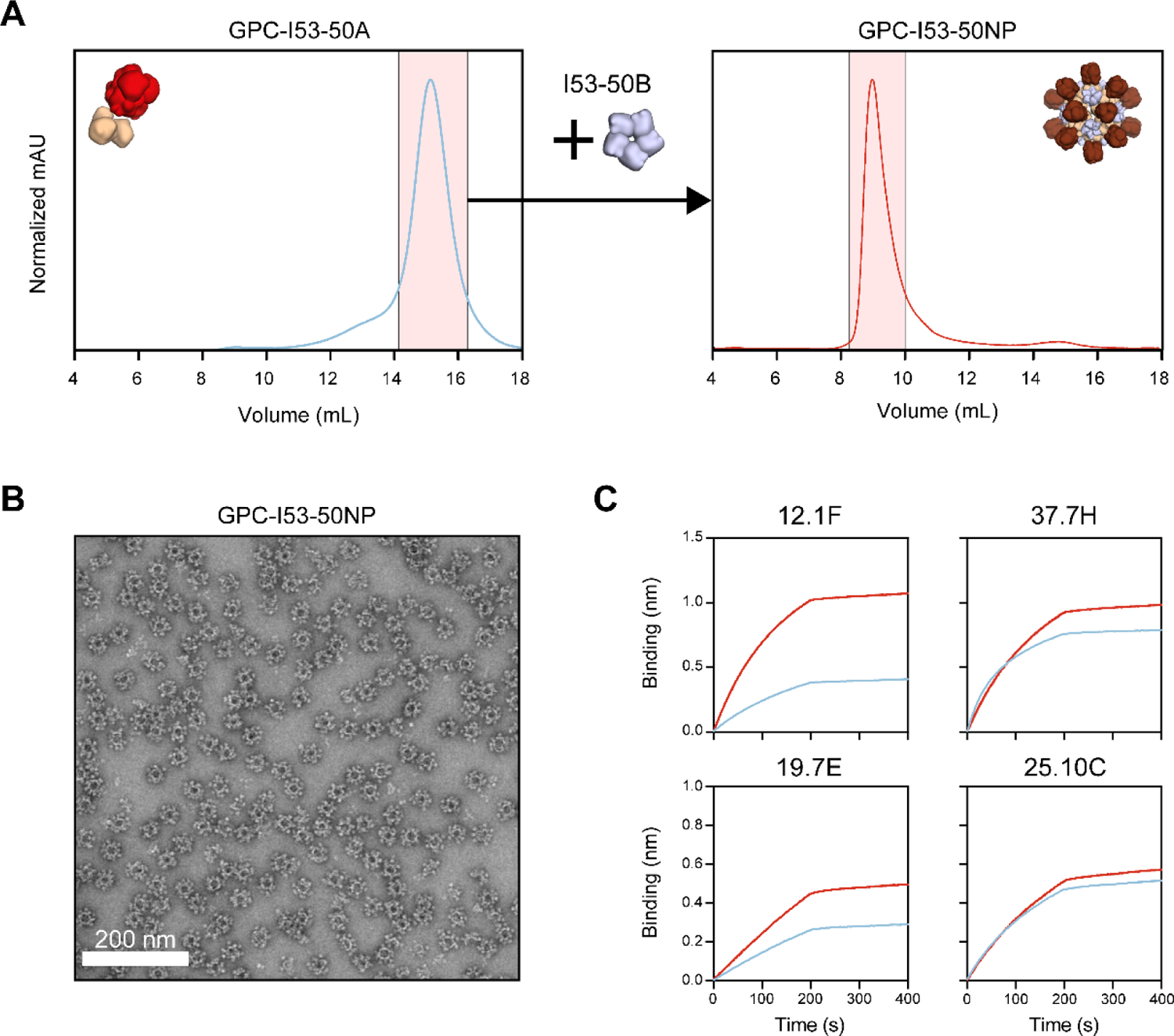
Biophysical and antigenic characterization of GPC-I53-50NPs. (A) Representative size exclusion chromatograph of GPC-I53-50A (left panel) and GPC-I53-50NPs after assembly with I53-50B (right panel). Collected fractions for particle assembly or of purified GPC-I53-50NPs are shown in pink shading. (B) Raw nsEM image of the SEC-purified GPC-I53-50NPs. White scale bar corresponds to 200 nm. (C) Sensorgrams from BLI experiments with GPC-I53-50A and GPC-I53-50NP showing the binding of bNAbs 12.1F, 37.7H, 19.7E, and 25.10C.

### High-resolution structure of unliganded LASV GPC reveals subtle structural differences to liganded LASV GPC

Next, the assembled GPC-I53-50NPs were subjected to structural characterization using cryoEM (see Table S1 and fig. S2 for details on data collection and processing). Due to the flexible linkage between the GPC and I53-50A, we were unable to reconstruct a single high resolution model of the entire nanoparticle antigen (Fig. 3A; note the lack of structural features on the GPC antigens). Therefore, we separately processed the signal originating from two flexibly linked entities, the GPC trimers and I53-50NP, resulting in 3D maps at 3.97 Å and 3.67 Å resolution, respectively (Fig. 3A). Structural models were relaxed into the reconstructed maps and model to map fits are shown (Fig. 3A, bottom row). The model refinement statistics are presented in Table S2.

**Fig. 3.**
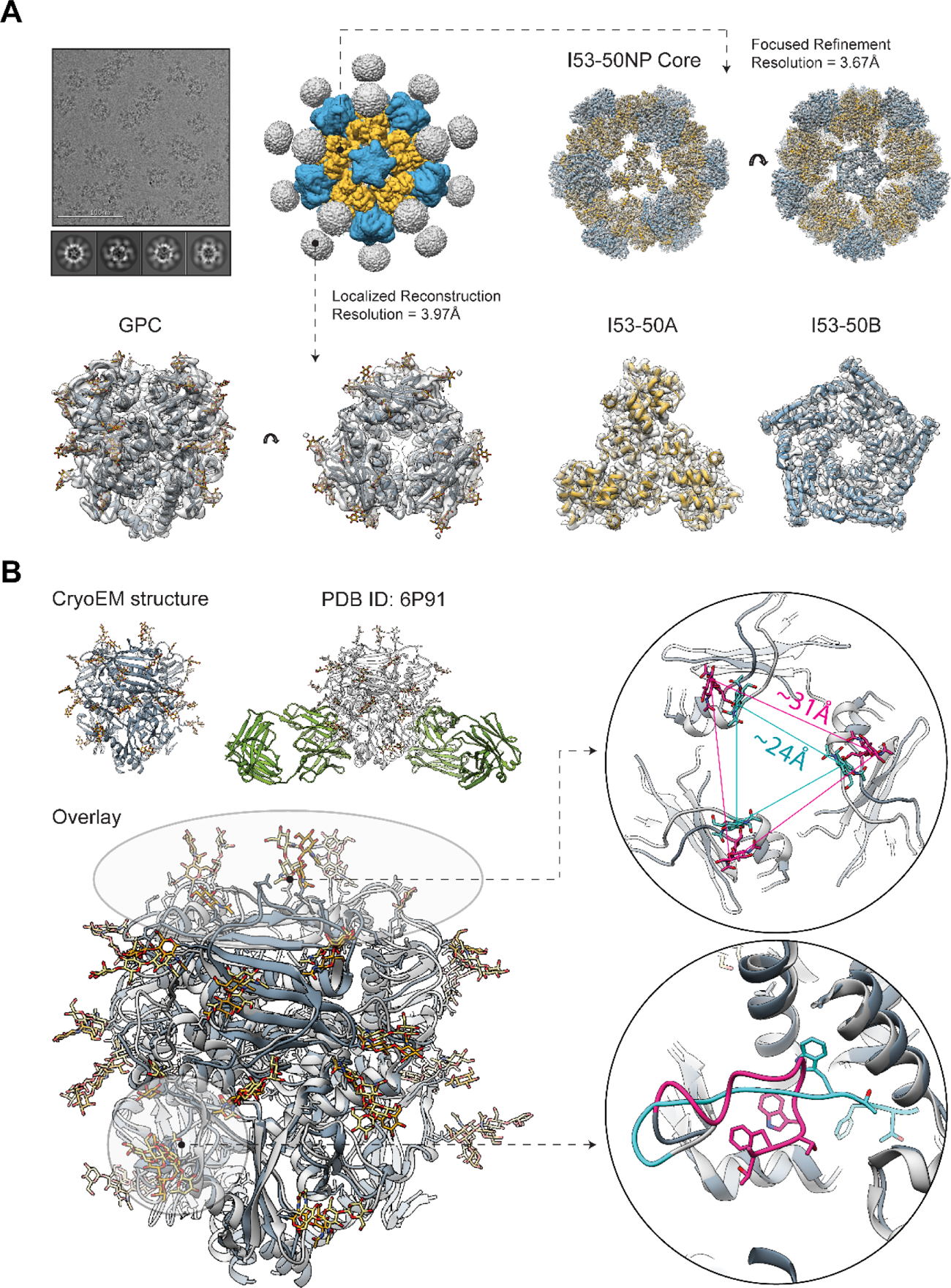
High-resolution cryo-EM structure of GPC-I53-50NPs. (A) Sample micrograph, 2D class averages and initial 3D reconstruction of the GPC-I53-50NP are displayed in the top left part of the panel. Focused refinement was applied to reconstruct the 3D map of the nanoparticle core (right). The structure of the I53-50NP is shown inside the map (I53-50A - yellow; I53-50B - blue; map - transparent white surface). Localized reconstruction approach was used for analysis of the presented antigen (bottom left). Refined GPC model is shown in dark gray with glycans displayed in golden yellow. (B) Comparison of the cryoEM structure of GPC (dark gray) and the crystal structure of GPC (light gray) in complex with 18.5C antibody (green) (PDB ID: 6P91; Hastie et al., 2019) with the overlay of the two structures shown below. Comparison of the fusion peptide conformations is displayed in the bottom right part of this panel (cryoEM model - pink; crystal structure - turquoise). Side-chains of residues G260 - W264 are displayed in each case. Comparison of the apex conformations in two structures is displayed in the top right part of this panel. The distances between N119 glycans are shown for each model (cryoEM model - pink; crystal structure - turquoise).

The I53-50NP structure is in excellent agreement with the Rosetta-designed model and the published structure obtained with I53-50NP presenting HIV-1 BG505 SOSIP trimers (PDB ID: 6P6F; Brouwer et al., 2019). The RMSD values for Cα atoms were 0.74 Å for the comparison with the Rosetta model and 0.51 Å for the comparison with HIV-1 SOSIP-I53-50NPs. These findings confirm that GPC fusion did not interfere with the folding of the I53-50A component or the assembly of the nanoparticle.

The reconstructed GPC model features a trimer in the prefusion state (Fig. 3A). The structure revealed an overall globular protein assembly, uniformly covered in N-linked glycans. In the cryoEM map we observed densities corresponding to all 11 N-linked glycans for each monomer. We compared this structure to the previously reported structures obtained by x-ray crystallography and featuring stabilizing antibodies bound to a quaternary epitope at the interface of two protomers, making direct contacts with the fusion peptide (Fig. 3B). Due to the high degree of similarity of the three crystal structures (PDB IDs: 6P91, 6P95 and 5VK2; (Hastie et al., 2017, 2019)) we are only showing the comparative analysis results for one of them (PDB ID: 6P91; (Hastie et al., 2019)). Overall, our unliganded cryoEM model displayed excellent structural homology with the reported antibody-complexed structure, with a Cα RMSD value of 0.92 Å suggesting that the fusion to the I53-50A component and nanoparticle assembly did not induce major conformational changes within the GPC.

Closer inspection of the two models revealed local differences in two regions: the fusion peptide and the trimer apex (Fig. 3B, bottom right). In the unliganded state (cryoEM structure; this study), fusion peptide residues G260-T274 expand into the cavity between two protomers with the N-terminus (residue G260) facing outwards. In the antibody-complexed states (PDB IDs: 6P91, 6P95 and 5VK2; (Hastie et al., 2017, 2019)) the cavity is occupied by the HCDR3 of the 18.5C, 25.6A or 37.7H antibodies, pushing the fusion peptide towards the center of the trimer. Solvent exposure of the fusion peptide in the unliganded prefusion state is a property shared by several other class 1 glycoproteins (e.g. influenza HA, and HIV Env) and may be important for the correct onset of the fusogenic conformational change in the GPC (White et al., 2008). The trimer apex also exhibited differences between the different structures (Fig. 3B, top right). The unliganded structure featured a more open conformation at the apex, particularly for residues between N114 and L128. The distances between the three N119 residues from the respective protomers is ∼31 Å in the unliganded structure and ∼24 Å in the antibody-complexed structures. This ∼7 Å difference is substantial and might affect the position and/or orientation of the glycans attached to the three N119 residues thereby influencing epitope shielding by these glycans. Finally, the local resolution of the unliganded model is lower within the central cavity of the trimer and at the apex (Fig. S2), suggesting a higher degree of flexibility in these regions. While the presence of stabilizing antibodies (18.5C, 25.6A or 37.7H) and crystal packing in the antibody-complexed structure may have resulted in increased stability of this part of the protein and more compact conformation, the current structural data is insufficient to conclude whether this conformational plasticity has any functional or immunological relevance.

### LASV GPC-I53-50NPs induce strong antibody responses in rabbits

To assess the immunogenicity of recombinant native-like GPC trimers and evaluate whether presentation on I53-50NPs would provide an immunological benefit, we performed an immunization study in rabbits with GPC-I53-50A and GPC-I53-50NPs. New Zealand White Rabbits (n = 6 per group) received 30 μg of GPC-I53-50A or an equimolar amount assembled into GPC-I53-50NPs formulated in squalene emulsion at weeks 0, 4, 16, and 28 (Fig. 4A). Bleeds were taken after the prime (week 4), and each boost (week 6, 18 and 30) to determine serum GPC-specific binding and pseudovirus neutralization titers.

**Fig. 4.**
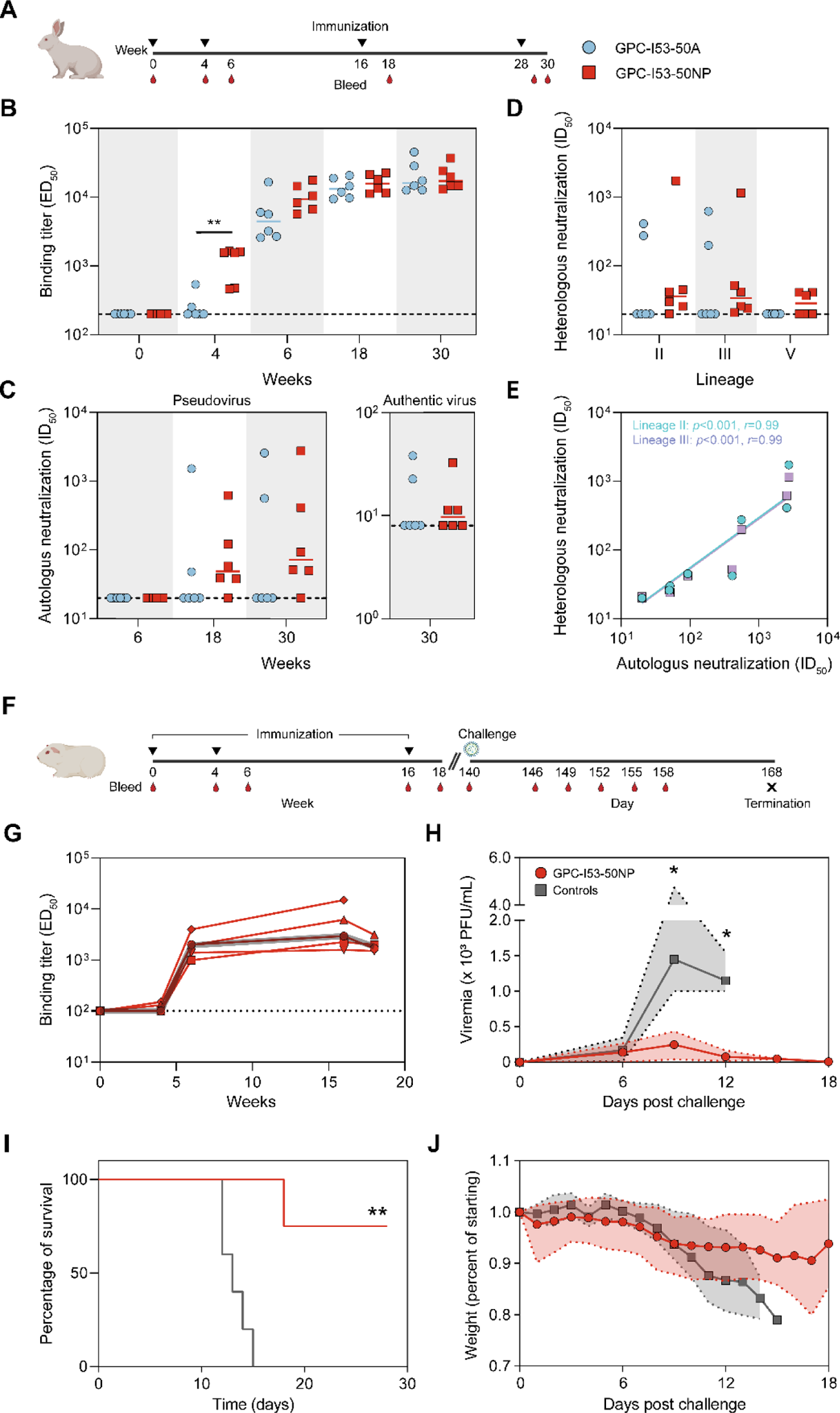
Immunogenicity of GPC-I53-50A and GPC-I53-50NP in rabbits and protective efficacy of GPC-I53-50NP in guinea pigs. (A) Schematic representation of the rabbit immunization schedule with color coding for each immunogen. (B) Midpoint binding titers against GPC-dn5B at week 0, 4, 6, 18, and 30. Horizontal bars represent medians. Statistical differences between two groups (*n* = 6 rabbits) were determined using two-tailed Mann–Whitney U-tests (\*\**p* < 0.01). (C) Midpoint NAb titers against autologous pseudovirus (lineage IV) at week 0, 4, 6, 18, and 30 (left panel). Endpoint NAb titers against authentic LASV (lineage IV) at week 30 (right panel). Horizontal bars represent medians. The dotted line indicates the lower limit of detection. (D) Midpoint NAb titers against heterologous pseudoviruses (lineage II, III and V) at week 30. Horizontal bars represent medians. (E) Correlation plot of autologous NAb titers vs heterologous NAb titers (lineage II or III). The *r* and *p* values are shown for non-parametric Spearman correlations (*n* = 7 rabbits; all rabbits with an autologous neutralization ID_50_ > 20). (F) Schematic representation of the guinea pig challenge study schedule. (G) Midpoint binding titers against GPC-dn5B at week 0, 4, 6, 16, and 18. The dark grey line represents the median. (H) Median RNA viral loads in vaccinated and control guinea pigs after challenge. The shaded area indicates the range. Statistical differences between two groups (week 9: *n* =4 for vaccinated, *n* = 5 for controls; week 12: *n* = 4 for vaccinated and controls) were determined using two-tailed Mann–Whitney U-tests (\**p* < 0.05). (I) Kaplan Meier-curve showing survival over time for vaccinated and control guinea pigs following a LASV challenge. Statistical difference between the two groups was determined using log-rank tests (\*\**p* < 0.005). The same color coding was used as in panel H. (J) Median weight alteration in vaccinated and control guinea pigs over time after challenge. The shaded area indicates the range. The same color coding was used as in panel H.

GPC-specific binding titers were analyzed by ELISA using immobilized GPC fused to the previously described dn5B scaffold to exclude serum Ab binding to I53-50A (Fig. 4B). Importantly, GPC-dn5B was almost exclusively trimeric as determined by nsEM and bound efficiently to 25.10C, 12.1F, 37.7H and 19.7E (Fig. S3). After the first immunization GPC-I53-50NPs induced significantly higher GPC binding titers than its soluble counterpart, which did not induce Ab titers above cutoff in 4/6 rabbits (median ED_50_ of 1557 versus 200, respectively; two-tailed Mann-Whitney *U*-test; *p*=0.0087). A trend towards higher binding titers was also observed after the second immunization, but after the third and fourth, GPC-specific binding titers were very similar between GPC-I53-50A and GPC-I53-50NP. We used a previously described lentivirus-based pseudovirus assay to measure the induction of NAb responses (Robinson et al., 2016). No neutralization was induced after two immunizations (Table S3). However, after three immunizations 5/6 rabbits in the nanoparticle group developed NAb titers with ID_50_ values ranging from 38 to 623. For the two best responders (rabbit 194 and 196) these titers increased to 2749 and 413 after the final fourth immunization, respectively (Table S3). In contrast, only 2/6 rabbits in the GPC-I53-50A group developed NAb responses after three immunizations which were boosted to titers of 2565 and 560 after the final boost (Fig. 4C and Table S3). In addition, we assessed the sera’s ability to neutralize authentic LASV (Fig. 4C and Table S4). In this assay the virus neutralization titer is calculated as the geometric mean titers (GMT) of the reciprocal value of the last serum dilution at which inhibition of the cytopathic effect on infected Vero E6 cells is detectable. Titers ranged from 11 to 38 in the rabbit sera that showed some neutralizing activity (2/6 rabbits in the GPC-I53-50A group and 3/6 in the GPC-I53-50NP group). Consistent with earlier studies, it is apparent that the authentic virus is considerably more resistant to neutralization than the lentivirus-based pseudovirus assay (Robinson et al., 2016).

Considering the high degree of sequence diversity of LASV lineages over the large geographical endemic areas, the ability to elicit a broad immunological response is imperative for a successful LASV vaccine. To evaluate the breadth of the NAb response induced by our vaccine based on a LASV lineage IV strain, we generated pseudoviruses representing the heterologous LASV lineages II, III and V and performed neutralization assays with rabbit serum from week 30 (two weeks after the fourth vaccination) (Fig. 4D and Table S3). Sera from the two rabbits that developed autologous NAb titers in the GPC-I53-50A group were also able to neutralize lineage II and III quite potently but not lineage V pseudovirus. Similarly, in the nanoparticle group, sera that neutralized lineage IV also neutralized lineage II and III to varying degrees. Serum of rabbit 194 showed broad and exceptionally high NAb titers to lineage II (ID_50_ of 1733) and III (ID_50_ of 1151) and low but detectable neutralizing activity to lineage V. We noted a strong correlation between NAb titers against autologous and heterologous (lineage II and III) pseudovirus, suggesting that our GPC immunogens readily elicit NAb against epitopes thar are conserved among LASV lineages (Fig. 4E).

### LASV GPC-I53-50NP vaccination protects guinea pigs from lethal LASV challenge

To assess the protective potential of GPC-I53-50NPs, we performed a LASV challenge study. Hartley guinea pigs (n = 5) received 30 μg of GPC-I53-50NP adjuvanted in squalene emulsion at weeks 0, 4, and 16 (Fig. 4F). Control animals (n = 5) received three doses of squalene emulsion. GPC-I53-50NPs induced GPC-specific Ab responses after two immunizations (median ED_50_ of 1941 at week 6) which did not increase any further after the third immunization (median ED_50_ of 1836 at week 18) (Fig. 4G). These GPC-specific Ab responses seem markedly lower than those observed in rabbits. In contrast to our observations in rabbits, no pseudovirus neutralization was detected after three immunizations with GPC-I53-50NPs in guinea pigs (Table S5). Together, this indicates that guinea pigs elicited an overall poorer humoral response than rabbits.

At week 20, i.e. four weeks after the third vaccination, the animals were challenged intraperitoneally with a lethal dose (1×10^4^ plaque-forming units (PFU)) of a guinea-pig adapted LASV Josiah strain (lineage IV; (Safronetz et al., 2015)) (Fig. 4F). One guinea pig in the vaccinated group did not recover from the week 18 bleed and died pre-challenge. With a median of 145 and 170 PFU/mL, both vaccinated and unvaccinated guinea pigs, respectively, had similar viral loads 6 days after the challenge. However, at day 9, viral loads had increased dramatically in the control animals reaching a median viral load of 1450 PFU/mL, which then decreased slightly to 1150 PFU/mL at day 12 after challenge. In contrast, the median viral loads in the vaccinated guinea pigs were significantly lower at day 9 (258 versus 1450 PFU/mL, two-tailed Mann-Whitney *U*-test; p = 0.0159) and day 12 (63 versus 1150; two-tailed Mann-Whitney *U*-test; p = 0.0286) (Fig. 4H). By day 15 all (5/5) control animals had met the euthanization criteria (Table S6) and were euthanized, while in the vaccinated guinea pigs viral loads had decreased further to a median of 38 PFU/mL. Nevertheless, on day 18 post-challenge, one of the vaccinated guinea pigs was moribund and had to be euthanized (Fig. 4I). The remaining three animals survived to the study endpoint (day 28). All GPC-I53-50NP-vaccinated animals became febrile after challenge and all but one experienced weight loss (Fig. 4J and fig. S4B). Thus, GPC-I53-50NP vaccination did not protect the animals from infection and disease, but it did significantly protect them from mortality (log-rank test; p = 0.0045).

### Isolation of a broadly neutralizing LASV GPC-specific mAb reveals a novel site of vulnerability

Most neutralizing mAbs isolated from Lassa fever patients so far reveal GPC-B, a base-proximate epitope that includes glycans at N390 and N395, as an immunodominant site for neutralization (Hastie et al., 2019; Robinson et al., 2016). Removal of these glycans drastically increased neutralization sensitivity of LASV to these NAbs, consistent with earlier reports showing that glycans are major barriers for the induction of potent NAb responses (Sommerstein et al., 2015; Watanabe et al., 2018). To assess if GPC-B-type NAbs dominated the induced NAb responses by GPC-I53-50A and GPC-I53-50NP in rabbits we performed neutralization assays using pseudovirus lacking PNGs at either glycan N390 or N395 (Fig. S5A). We observed no clear alteration in NAb titers when these glycans were removed suggesting that GPC-B-type NAbs did not dominate the neutralizing responses induced by GPC-I53-50NPs.

To characterize the NAb response in rabbit 194 in more detail we sorted B cells and isolated mAbs. Rabbit 194 was immunized with GPC-I53-50NPs and showed the most potent and broad neutralization profile. GPC-specific IgG+ B cells were single-cell sorted from peripheral blood mononuclear cells (PBMCs) by dual staining with fluorescently labeled GPC-I53-50A from the Josiah (lineage IV) and NIG08-A41 (lineage II) strain. Of the GPC-specific B cells (∼15.5% of the total IgG+ B cells), ∼93% was reactive to both probes, consistent with the breadth of the NAb response in this animal (Fig. 5A). We cloned four mAbs, termed LAVA01-LAVA04, which bound to GPC-dn5B with a midpoint binding concentration (EC_50_) varying from 0.02 to 0.10 µg/mL (Fig. S5B). LAVA01 was able to neutralize autologous pseudovirus with an IC_50_ of approximately 0.12 µg/mL; on par with that of 19.7E but approximately 10-fold lower than 37.7H (Fig. 5B and S5C). LAVA01 also showed neutralization potency to NIG08-A41 (lineage II) and CSF (lineage III) pseudovirus with an IC_50_ of 0.78 µg/mL and 1.08 µg/mL, respectively (Fig. 5C). In contrast, the prototypic bNAb 37.7H failed to reach 50% neutralization of CSF. Neither of the three mAbs, LAVA01, 37.7H, or 19.7E, were able to reach 50% neutralization of the Bamba strain (lineage V) (Fig. S5D). We also analysed the potential of LAVA01 to neutralize authentic Josiah LASV. LAVA01 neutralized authentic LASV at a titer of 21 µg/mL (Fig. 5D). 19.7E was less potent than LAVA01 showing a neutralization titer of 42 µg/mL, whereas 37,7H was more potent, neutralizing authentic LASV with a titer of 5 µg/mL.

**Fig. 5.**
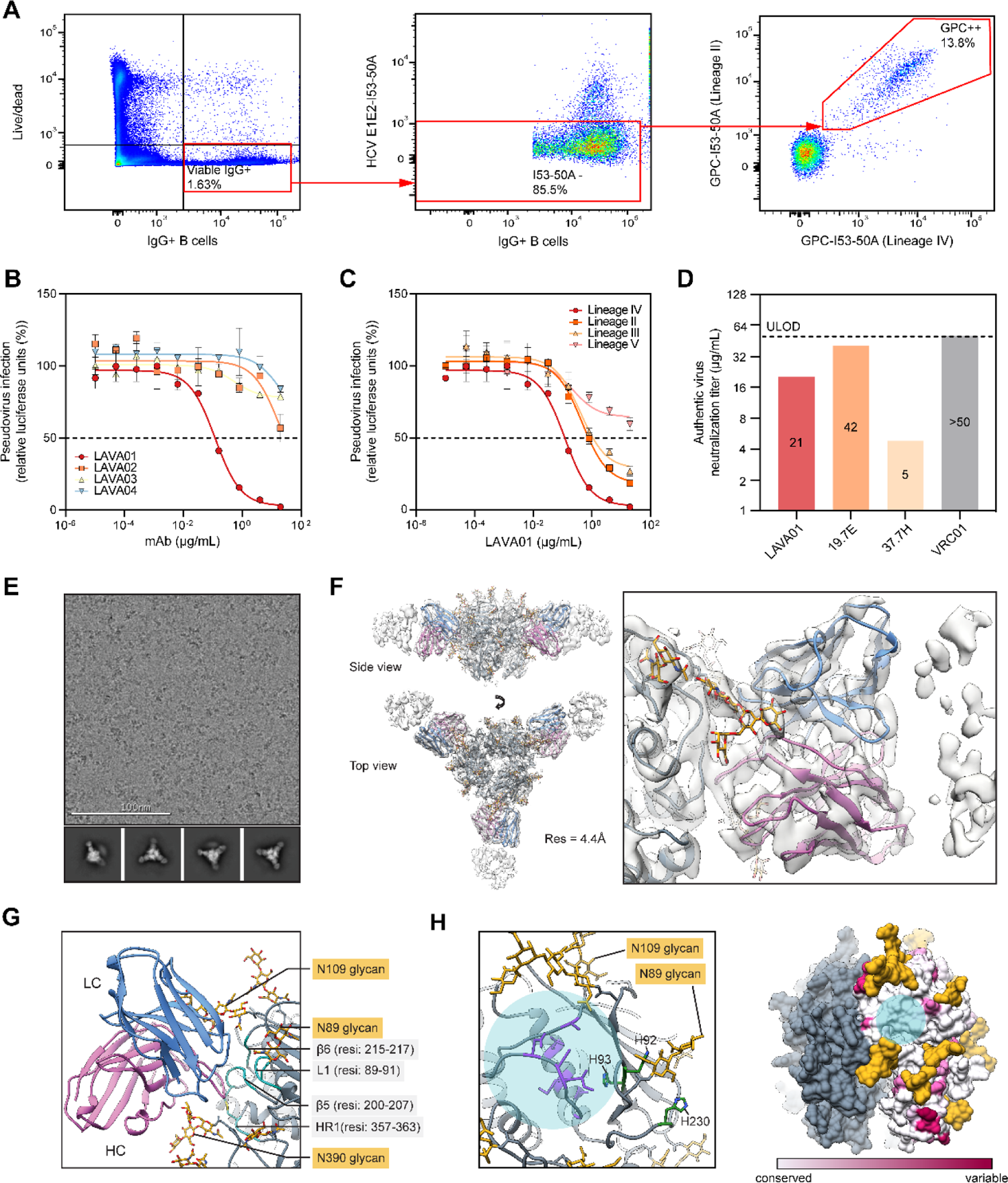
Isolation of a monoclonal NAb that targets a novel site of vulnerability on GPC. (A) Representative gating strategy for the isolation of GPC-specific B cells. The lymphocyte population was selected and doublets were excluded (not shown). From live IgG+ B cells (left), cells were selected that showed low reactivity to I53-50A (middle) after which the double-positive GPC-specific B cells were sorted (right). (B) Pseudovirus neutralization of Lineage IV (Josiah) by LAVA01-LAVA04. The dotted line indicates 50% neutralization. Shown are the mean and SEM of two technical replicates. Symbols are the same as indicated in panel B. (C) Pseudovirus neutralization of Lineage II, III, IV and V by LAVA01. The dotted line indicates 50% neutralization. Shown are the mean and SEM of two technical replicates. (D) Endpoint neutralization titers against authentic LASV (lineage IV) by LAVA01, 19.7E, 37.7H and VRC01 (HIV-1 Env-specific mAb; negative control). The dotted line indicates the upper limit of detection. (E) Representative raw micrograph and 2D class averages of LAVA01 Fab complexed with chemically cross-linked GPC-I53-50A (F) Reconstructed 3D map (transparent white surface) and relaxed model (GPC - gray, glycans - golden, LAVA01 heavy chain - purple, LAVA01 light chain - blue) shown in top and side view with a close-up view of the antibody-antigen interface. (G) Peptide (turquoise) and glycan (golden) epitope components for LAVA01 antibody. (H) Overlay of the LAVA01 footprint (transparent light blue circle) with the histidine triad (green) and proposed LAMP-1 receptor binding site residues determined by mutagenesis (purple; left panel). Sequence conservation of GPC with the LAVA01 footprint shown (transparent light blue circle; right panel).

To map the epitope of LAVA01 we performed neutralization assays with pseudovirus (lineage IV) that had the N390 or N395 glycan knocked out. Whereas removing the N395 glycan slightly decreased LAVA01 neutralization, removal of the N390 glycan completely abrogated it, suggesting that the N390 glycan is a key component of the LAVA01 epitope (Fig. S5E).

To obtain detailed structural information of the epitope of this new NAb we performed cryo-EM studies using GPC-I53-50A in complex with the LAVA01 Fab. Initial attempts revealed that the large majority of the trimers disassembled into monomers upon complex formation, resulting in highly heterogeneous 2D-classes which could not be used to reconstruct a 3D-map (Fig. S6). To resolve this problem we chemically cross-linked GPC-I53-50A with glutaraldehyde prior to complexing with the LAVA01 Fab. This increased the relative amount of trimeric GPC; although monomers were still visible in the micrographs and 2D classes (Fig. 5E and S6). Data processing resulted in a 4.41 Å map of the complex, from which we derived an atomic model (Fig. 5F). Due to the relatively modest resolution, amino-acid side-chains of the LAVA01 Fab were not built past Cβ. Inspection of the model showed that LAVA01 binds to a previously unknown neutralizing epitope comprising elements of GP1 and GP2 within a single protomer and can bind the GPC trimer at the maximum stoichiometry of 3. The main peptidic components of the epitope are β5, β6, L1 and HR1 (Fig. 5G). Moreover, the antibody makes extensive contacts with the N109 glycan, through the HCDR2 and LCDR1 loops, and with the N390 glycan, through the HCDR2 and HCDR3 loops. The latter explains the strong glycan-dependent neutralization of LAVA01. Even though side chains were not explicitly built, the high amount of tyrosine residues were readily discernible on the LAVA01 heavy chain, mediating interactions with glycan residues, similarly to several HIV-1 bNAbs (Lee et al., 2017; McLellan et al., 2011). The location of the epitope provides the basis for a potential neutralization mechanism for LAVA01. First, this site is in immediate proximity to the histidine triad consisting of H92, H93 and H230 that is proposed to regulate the onset of the pH-induced conformational change in GP1 that precedes LAMP-1 binding and membrane fusion. Second, mutagenesis studies have shown identified β5 residues 200-207 as being critical for LAMP-1 receptor binding (Israeli et al., 2017) (Fig. 5H, left). We thus propose that LAVA01 inhibits LASV by interfering with the membrane fusion step through disruption of the fusogenic conformational change at low pH and/or by sterically blocking the binding to LAMP-1. Notably, this epitope features a relatively high sequence conservation across known LASV strains (Fig. 5H, right), consistent with the critical role it has in viral entry. These data are in agreement with the pseudovirus neutralization experiments described above where we observed cross-neutralization of LAVA01 with different lineages of LASV. Collectively, neutralization assays and cryo-EM work showed that we have isolated a relatively broad neutralizing antibody, elicited after vaccination, that targets a novel site of vulnerability on LASV GPC.

## Discussion

Even though the generation of a prefusion stabilized GPC represents an important advancement for the development of a recombinant protein vaccine, its propensity to dissociate into monomers compromises its potential to induce potent NAb responses (Hastie et al., 2017, 2019). The majority of the NAb epitopes require a trimeric GPC, while the exposed interior of monomeric GPCs presumably constitute a highly immunogenic non-neutralizing epitope as it is the only surface that is not covered by glycans. So far, the detailed in vitro and in vivo characterization of a soluble prefusion-stabilized trimeric GPC immunogen have not been reported. Here, we showed that fusing GPC to I53-50A facilitates trimer stabilization and enables the formation of nanoparticles presenting native-like trimers. This work allowed us to acquire an antibody-free high-resolution structure of LASV GPC while potentially generating a pipeline for solving GPC structures from other LASV lineages as well as antibody-complexed structures. An immunogenicity study in rabbits demonstrated that GPC-I53-50NPs can induce strong humoral immune responses, including neutralizing responses. In addition, we showed that these nanoparticle immunogens can protect guinea pigs from LASV-induced mortality. Furthermore, isolation of mAbs from immunized rabbits revealed that these nanoparticles are able to elicit antibody responses with broad neutralizing potency that target a previously unreported epitope on the putative LAMP-1 binding site.

The absence of the trimer-stabilizing helical bundle that is positioned in the center of most viral type 1 fusion glycoproteins, as well as the large cavities in the trimeric interphase, likely explain the intrinsic instability of recombinantly expressed prefusion GPC trimers (Hastie and Saphire, 2018; Hastie et al., 2017). Fusion to I53-50A did not strengthen the relatively weak interactions between the GPC protomers but functioned essentially as a trimerization domain. This allowed GPC to be expressed at feasible yields while improving antigenicity. Even though electron microscopy studies clearly demonstrated that fusion to I53-50A facilitates the generation of native-like GPC trimers, there is still room for improvement. Splayed open GPC trimers kept together by I53-50A were visible in the 2D-class averages, which became more abundant when complexed with LAVA01 Fab. Additional stabilization strategies might be necessary to further improve trimerization. We showed here that chemically cross-linking GPC-I53-50A is a promising avenue. Alternatively or concomitantly, shortening or rigidifying the linker between GPC and the trimeric scaffold may represent straightforward approaches to further stabilize trimers. Indeed, GPC-dn5B, which we generated during the course of this study for serum ELISAs and contained only a 3-amino acid linker, showed a much higher trimeric homogeneity than GPC-I53-50A by nsEM.

Three immunizations with GPC-I53-50A were necessary to induce pseudovirus neutralization in a subset of animals, which increased to potent NAb titers after the final and fourth boost. Presentation on I53-50NPs improved GPC’s ability to induce NAb responses with only 1 rabbit showing no neutralization after four immunizations. In addition, we also observed that GPC-I53-50NPs induced heterologous neutralization to a lineage II and III pseudovirus, even in the context of low autologous NAb titers. Moreover, sera from GPC-I53-50NP recipients, in contrast to those from animals immunized with trimer only, were able to neutralize a lineage V pseudovirus albeit with low potency. Consistent with our study, four immunizations with a LASV GPC-VLP were required to induce potent and broad neutralization, with NAb titers being the lowest to lineage V (Müller et al., 2020). However, despite potent pseudovirus neutralization in some animals, authentic LASV neutralization was considerably weaker. The large difference in sensitivity between the two assays stresses the importance of including assays that use authentic LASV when assessing NAb responses. The absence of a NAb response in the immunized guinea pigs was surprising. This may relate to the overall weaker responses, differences in the B cell repertoire, or to other factors. Further stabilization of GPC, by aforementioned or alternative trimer engineering strategies, may improve the immunogenicity of these nanoparticle immunogens. Increasing the overall immunogenicity of GPC nanoparticles might also be necessary.

Monoclonal Ab isolation yielded LAVA01, which neutralized lineage IV pseudovirus as well as lineage II and III, the two lineages which currently dominate LASV outbreaks in Nigeria (Kafetzopoulou et al., 2019). The neutralization potency against these pseudoviruses was similar to that of the GP1-targeting human NAb 19.7E. Interestingly, the bNAb 37.7H was unable to neutralize lineage III pseudovirus in our hands. This is in contrast to previous results with the same pseudovirus assay (Robinson et al., 2016), but consistent with other reports (Müller et al., 2020). Furthermore, whereas 19.7E was previously unable to neutralize lineage III (Robinson et al., 2016), it did neutralize our lineage III pseudovirus. These results may be explained by the sequence variation between the different lineage III strains used and highlight the need for a standardized panel of reference strains for assessment of neutralization breath. The relatively broad neutralizing phenotype of LAVA01 may be the result of this mAb targeting highly conserved residues on the LAMP-1 binding site and making extensive contacts with the conserved glycans at sites N109 and N390. The dependency of pseudovirus neutralization on glycan N390 is an interesting feature as it reveals that the glycan shield may not only shield GPC from NAb responses but can also be a target, akin to the glycan shield of HIV-1 Env (Seabright et al., 2019, 2020). The epitope of LAVA01 represents a new target to be exploited by vaccines and antibody therapeutics.

The relevance of NAbs as a protective endpoint for LASV vaccines has only recently been made clear through the success of several passive immunization studies with monoclonal bNAbs in animal models (Cross et al., 2016, 2019; Mire et al., 2017). Yet, various studies have also shown protection to LASV challenge in the absence of measurable (pseudovirus) NAb responses, suggesting a role for cellular immunity or non-neutralizing Ab effector functions in vaccine-induced protection (Abreu-Mota et al., 2018; Fischer et al., 2021; Mateo et al., 2021). Our guinea pig study provides additional evidence that NAb responses may not be the sole determinant for protection. However, GPC-I53-50NP’s inability to prevent illness and fully protect against a lethal LASV challenge in guinea pigs suggest that GPC-I53-50NP’s efficacy may be improved when able to induce a potent NAb response. This study represents a step forward in generating a LASV vaccine that induces strong and protective humoral immunity.

## Acknowledgments

We thank Robin Shattock and Paul McKay at Imperial College London for kindly sharing the full-length lineage IV (Josiah) GPC vector, Dietmar Katingen and Philipp Mundsperger at Polymun Scientific for providing the SE adjuvant, and Rashmi Ravichandran at the Institute for Protein Design for sharing purified I53-50B. We thank Gotthard Ludwig and Sebastian Schmidt from the biosafety level 4 facility at the Philipps-University of Marburg for technical assistance. The authors also thank Bill Anderson and Hannah Turner from The Scripps Research Institute for their help with electron microscopy experiments. The rabbit image in figure 4 was created with BioRender.com. This work was supported by a Vici fellowship from the Netherlands Organisation for Scientific Research (NWO; to R.W.S.), by the Fondation Dormeur, Vaduz (to R.W.S. and to M.J.v.G.), and the Bill and Melinda Gates Foundation through grant OPP1170236 (to A.B.W.). A.A. is supported by the amfAR Mathilde Krim Fellowship in Biomedical Research (#110182-69-RKVA). T.S. received funding from the Deutsche Forschungsgemeinschaft (DFG, German Research Foundation)—Projektnummer 197785619/SFB1021.

## Author Contributions

P.J.M.B., A.A., A.J.R., H.M., T.Str., A.B., A.B.W., and R.W.S. conceived and designed experiments. P.J.M.B., A.A., A.J.R., H.M., Y.W., M.Cl., N.M.L., T.P.L.B., H.R.P., T.Ste., and J.A.B. performed the experiments. P.J.M.B., A.A., A.J.R., H.M., Y.W., M.Cl., M.M.H., and M.J.G set up experimental assays. P.J.M.B., A.A., A.J.R., H.M., Y.W., M.Cl., A.B.W., and R.W.S. analyzed and interpreted data. K.S. shared the HCV E1E2-I53-50A reagent for FACS sorting. P.J.M.B., A.A., A.J.R., H.M., Y.W., T.Str., A.B.W., and R.W.S. wrote the manuscript with input from all authors.

## Data availability

3D maps from electron microscopy experiments have been deposited to the Electron Microscopy Databank (http://www.emdatabank.org/) under accession IDs: EMD-25107, EMD-25108, EMD-25109. 3D models from electron microscopy experiments have been deposited to the Protein Data Bank (http://www.rcsb.org/) under accession IDs: 7SGD, 7SGE, 7SGF.

## Method Details

### Construct design

To generate the prefusion stabilized GPC construct, a gene encoding residues 1-424 of GPC from the Josiah strain (Genbank: ADY11068.1) with the GPCysR4 mutations (R207C, E329P, L258R, L259R, G360C; (Hastie et al., 2017)) following C-terminal extension, GSGSLEWSHPQFEK (GS encodes a BamHI site), was cloned into a PstI-NotI-digested pPPI4 plasmid by Gibson assembly. GPC-I53-50A was created by digesting the GPC plasmid with BamHI and NotI and subsequent Gibson assembly with a gene encoding the previously described I53-50A.1NT1 sequence (including the glycine-serine linker) with a GSLEWSHPQFEK extension on the C-terminus (Brouwer et al., 2021b). For the GPC-dn5B construct a gene encoding the recently described I53_dn5B sequence (EEAE…MREE; (Ueda et al., 2020)) preceded by a GSG sequence and followed by a GGWSHPQFEK sequence was ordered and cloned into a BamHI-NotI digested GPC-I53-50A plasmid. The lineage IV probe for B cell sorting was generated by Gibson assembly of the previously mentioned prefusion GPC gene into a PstI-BamHI-digested pPPI4 plasmid encoding a I53-50A.1NT1 sequence that had an Avi- and histidine-tag after the final residue. To generate a lineage II probe a gene encoding residues 1-423 of GPC from the NIG08-A41 strain (Genbank: ADU56626) containing the GPCysR4 mutations were introduced. Plasmids encoding the mAbs 19.7E, 37.7H, 12.1F, and 25.10C were generated by ordering genes encoding the variable regions of the corresponding heavy and light chains and cloning them in expression vectors containing the constant regions of the human IgG1 for the heavy or light chain using Gibson assembly. For pseudovirus neutralization assays, pPPI4 plasmid was digested with PstI-NotI and a gene encoding full-length GPC of lineage II (NIG08-A41), lineage III (CSF; Genbank: AAL13212.1), or lineage V (Bamba; AHC95555.1) was inserted by Gibson assembly. The N390D and N395D mutants were generated by Q5 site-directed mutagenesis using a plasmid encoding full-length Josiah strain as a template.

### Protein expression and purification

Proteins were expressed by transient transfection of HEK 293F cells (0.8-1.2 million cells/mL) maintained in Freestyle medium (Life Technologies). A 3:1 ratio of PEImax and expression plasmids (312.5 µg/L cells) were added to the cells. For GPC fusion proteins a 2:1 ratio of GPC-I53-50A/GPC-I53-dn5B and furin was used to ensure optimal furin cleavage. For non-scaffolded GPC this was 3:1. MAbs were transfected using a 1:1 ration of heavy and light chain expression plasmid. After six days, supernatants were harvested by centrifugation (30 min at 4000 rpm) and filtration using a 0.22 µm Steritop filter (Merck Millipore). Supernatants containing mAbs were purified as described previously (Brouwer et al., 2020). GPC constructs were purified using StrepTactinXT Superflow high capacity 50% suspension in accordance with the manufacturer’s protocol for gravity flow (IBA Life Sciences). Prior to loading on the column, Biolock solution and a 10X buffer W (1 M Tris/HCl, 1.5 M NaCl, 10 mM EDTA, pH 8.0) were diluted 1:1000 and 1:10, respectively, in the filtered supernatant. Eluted GPC constructs were buffer exchanged in Tris-buffer-saline (TBS), supplemented with 5% glycerol, using Vivaspin filters with a 50,000 kDa molecular weight cutoff (GE Healthcare). GPC-I53-50A for rabbit immunization studies were subjected to an additional SEC step using a Superose 6 increase 10/300 GL column in TBS, 5% glycerol. Fractions between 14 and 16.5 mL were collected, pooled and concentrated using Vivaspin filters with a 50,000 kDa molecular weight cutoff (GE Healthcare). The Nanodrop was used to determine the concentrations of expressed proteins applying the proteins peptidic molecular weight and extinction coefficient as calculated by the online ExPASy software (ProtParam).

### GPC-I53-50NP assembly

GPC-I53-50NPs were generated by collecting the previously mentioned SEC fractions of non-aggregated GPC-I53-50A (14 - 16.5 mL) and adding I53-50B.4PT1 pentamer (expressed as described previously (Walls et al., 2020)) at an equimolar amount of monomeric subunits. After an overnight incubation at 4°C, the mix was run over a Superose 6 increase 10/300 GL column in TBS, 5% glycerol to remove non-assembled components. Fractions between 8.5 and 10 mL were pooled and concentrated by centrifugation at 350 x *g* using Vivaspin filters with a 10,000 kDa molecular weight cutoff (GE Healthcare). Concentrated GPC-I53-50NPs were then diluted 1:1 with TBS, 5% glycerol, 400mM glycine, and the protein concentration was determined by Nanodrop method using the peptidic molecular weight and extinction coefficient as calculated by the online ExPASy software (ProtParam).

### BN-PAGE and SDS-PAGE analysis

BN-PAGE and SDS-PAGE were performed as described previously (de Taeye et al., 2015). Briefly, for BN-PAGE analysis, 2.5 μg of GPC or GPC-I53-50A was mixed with loading dye and run on a 4%–12% Bis-Tris NuPAGE gel (Invitrogen). For SDS-PAGE analysis, 2.5 ug of GPC-I53-50A or GPC-I53-50NP were mixed with loading buffer in the presence or absence of dithiothreitol and denatured, before loading on a 10-20% Tris-Glyine gel (Invitrogen).

### Negative stain electron microscopy

Negative stain electron microscopy experiments were performed as described previously (Antanasijevic et al., 2020). GPC, GPC-I53-50A and GPC-I53-50NP samples were diluted to 10-20 µg/ml, and 3 µl were loaded onto carbon-coated Cu grids (400-mesh). Prior to sample application the grids were glow-discharged at 15 mA for 25 s. Following a 10 s incubation period the samples were blotted off and the grids were negatively stained with 3 µl of 2% (w/v) uranyl-formate solution for 60 s. The grids were imaged on a Tecnai Spirit electron microscope (operating at 120 keV, nominal magnification was 52,000 X, resulting pixel size at the specimen plane was 2.05 Å). Electron dose was set to 25 e^-^/Å^2^ and the nominal defocus for imaging was −1.50 μm. Micrographs were recorded with a Tietz 4k x 4k TemCam-F416 CMOS camera. For data acquisition we used Leginon automated imaging software (Suloway et al., 2005). Appion data processing suite (Lander et al., 2009) was applied for all processing steps (particle picking, extraction and 2D classification).

### Bio-layer interferometry

The mAbs 19.7E, 12.1F, 37.7H or 25.10C diluted in running buffer (PBS, 0.02% Tween20, 0.1% BSA) were loaded on a Protein A sensor to a signal of 1.0 nm using an Octet K2 system (ForteBio). After removal of excess mAb by a short dip in running buffer the chip was dipped for 200 s in a concentration of 100 nM GPC, GPC-I53-50A or 5 nM of GPC-I53-50NP, diluted in running buffer. After this association step, the chip was dipped for 200 s in running buffer to measure protein dissociation. The same procedure was used for the comparison of GPC-I53-50A with GPC-dn5B except that Anti-Human Fc Capture (AHC) sensors were used, the sensors were loaded to a signal of 1.5 nm, and a 200 nM concentration of GPC-I53-50A or GPC-dn5B was used.

### Differential Scanning fluorimetry

Prometheus NT.48 NanoDSF instrument (NanoTemper Technologies) was used for the DSF experiments, as described previously (Antanasijevic et al., 2020). GPC, GPC-I53-50A and GPC-I53-50NP samples were diluted to 1 mg/ml in the TBS buffer (Alfa Aesar) and ∼10 µl of each diluted sample (in duplicates) was loaded into NanoDSF capillaries (NanoTemper Technologies). The temperature was raised from 20°C to 95°C at 1°C/min rate. The *T*_m_ value was determined from the first derivative curves using the NT.48 NanoDSF instruments software. The average value from the duplicate measurements is reported as the *T*_m_ value in the manuscript.

### Site-specific glycan analysis

Three 30 μg aliquots of GPC-I53-50A were denatured for 1h in 50 mM Tris/HCl, pH 8.0 containing 6 M of urea and 5 mM dithiothreitol (DTT). Next, the glycoprotein were reduced and alkylated by adding 20 mM iodoacetamide (IAA) and incubated for 1h in the dark, followed by a 1h incubation with 20 mM DTT to eliminate residual IAA. The alkylated glycoproteins were buffer-exchanged into 50 mM Tris/HCl, pH 8.0 using Vivaspin columns with a 3 kDa molecular weight cutoff and digested separately overnight using trypsin chymotrypsin or Glu-C (Mass Spectrometry Grade, Promega) at a ratio of 1:30 (w/w). The next day, the peptides were dried and extracted using C18 Zip-tip (Merck Milipore). The peptides were dried again, re-suspended in 0.1% formic acid and analyzed by nanoLC-ESI MS with an Easy-nLC 1200 (Thermo Fisher Scientific) system coupled to a Fusion mass spectrometer (Thermo Fisher Scientific) using higher energy collision-induced dissociation (HCD) fragmentation. Peptides were separated using an EasySpray PepMap RSLC C18 column (75 µm × 75 cm). A trapping column (PepMap 100 C18 3μM 75μM x 2cm) was used in line with the LC prior to separation with the analytical column. The LC conditions were as follows: 275 minute linear gradient consisting of 0-32% acetonitrile in 0.1% formic acid over 240 minutes followed by 35 minutes of 80%acetonitrile in 0.1% formic acid. The flow rate was set to 200 nL/min. The spray voltage was set to 2.7 kV and the temperature of the heated capillary was set to 40°C. The ion transfer tube temperature was set to 275°C. The scan range was 400−1600 m/z. The HCD collision energy was set to 50%, appropriate for fragmentation of glycopeptide ions. Precursor and fragment detection were performed using an Orbitrap at a resolution MS1= 100,000. MS2= 30,000. The AGC target for MS1=4e5 and MS2=5e4 and injection time: MS1=50ms MS2=54ms.

Glycopeptide fragmentation data were extracted from the raw file using ByonicTM (Version 3.5) and ByologicTM software (Version 3.5; Protein Metrics Inc.). The glycopeptide fragmentation data were evaluated manually for each glycopeptide; the peptide was scored as true-positive when the correct b and y fragment ions were observed along with oxonium ions corresponding to the glycan identified. The MS data was searched using the Protein Metrics N-glycan library. The relative amounts of each glycan at each site as well as the unoccupied proportion were determined by comparing the extracted chromatographic areas for different glycotypes with an identical peptide sequence. All charge states for a single glycopeptide were summed. The precursor mass tolerance was set at 4 ppm and 10 ppm for fragments. A 1% false discovery rate (FDR) was applied. The relative amounts of each glycan at each site as well as the unoccupied proportion were determined by comparing the extracted ion chromatographic areas for different glycopeptides with an identical peptide sequence. Glycans were categorized according to the composition detected. HexNAc(2)Hex(9−5) was classified as M9 to M5. HexNAc(3)Hex(5−6)(X) was classified as Hybrid with HexNAc(3)Fuc(1)(X) classified as Fhybrid. Complex-type glycans were classified according to the number of processed antenna and fucosylation, as HexNAc(3)(X), HexNAc(3)(F)(X), HexNAc(4)(X), HexNAc(4)(F)(X), HexNAc(5)(X), HexNAc(5)(F)(X), HexNAc(6+)(X), HexNAc(6+)(F)(X).

### Dynamic light scattering

DLS was used to assess the hydrodynamic radius (*R*_h_) and polydispersity of the assembled SOSIP-I53-50NPs. The particles were diluted to 0.025 µg mL−1 in PBS and loaded into a Dynapro Nanostar instrument (Wyatt Technology Corporation). *R*_h_ and polydispersity values were measured with ten acquisitions of 5 s each at 25°C and analyzed using the manufacturer’s software (Dynamics, Wyatt Technology Corporation), while assuming particles with a spherical shape.

### Preparation of LASV glycoprotein immune complex with LAVA01 Fab

To further stabilize the trimeric conformation, purified GPC-I53-50A was chemically cross-linked using glutaraldehyde. A concentration of 0.75 µg/ml GPC-I53-50A in PBS was mixed in a 1:1 volume ratio with PBS, 60 mM glutaraldehyde and incubated for 5 min at RT. The cross-linking reaction was then stopped by adding Tris, pH 7.4 to a final concentration of 75 mM, followed by an incubation step of 10 min at RT. Subsequently, GPC-I53-50A was dialysed twice to TBS and then twice to PBS using the Slide-A-Lyzer Mini dialysis devices with a 10,000 molecular weight cutoff (Thermo Fisher Scientific). Finally, GPC-I53-50A was concentrated to at least 2 µg/ml using Vivaspin centrifugal filters (GE Healthcare). To generate complexes, 400 µg of LAVA01 antibody (as Fab fragment) was incubated with 200 µg of cross-linked GPC-I53-50A trimers for 1 hour at room temperature. The complex was purified from excess/unassembled material using size-exclusion chromatography (HiLoad® 16/600 Superdex® pg200 (GE Healthcare) column running TBS buffer (Alfa Aesar)), and concentrated to 3 mg/ml using the Amicon ultrafiltration units with 10 kDa cutoff (EMD Millipore).

### CryoEM grid preparation and imaging

For the preparation of cryoEM grids the GPC-I53-50NP sample (1) and the GPC-I53-50A immune complex with LAVA01 Fab (2) were concentrated to 2 mg/ml and 3 mg/ml, respectively. Vitrobot mark IV was used for the preparation of cryoEM grids. The settings were as follows: temperature inside the chamber was 10 °C, humidity was 100%, blotting force was 0, wait time was 10 s, blotting time was varied within a 3-6 s range. Lauryl maltose neopentyl glycol (LMNG) detergent at a final concentration of 0.005 mM was added to each sample and 3 µl of that solution was immediately loaded onto plasma-cleaned QuantiFoil R 2/1 (400-mesh; Quantifoil Micro Tools GmbH) and UltrAuFoil R 1.2/1.3 grids (300-mesh; Quantifoil Micro Tools GmbH). The plasma cleaning step was performed in the Solarus 950 plasma system (Gatan) with Ar/O_2_ gas mix for 7 s. The sample was blotted off and the grids were plunge-frozen into liquid-nitrogen-cooled liquid ethane. Cryo-grids were imaged on an FEI Titan Krios (Thermo Fisher Scientific) microscope operating at 300 keV, equipped with a sample autoloader and the K2 Summit direct electron detector camera (Gatan). Exposure magnification was set to 29,000 and the resulting pixel size at the specimen plane was 1.03 Å. Automated image collection was performed using the Leginon software suite (Suloway et al., 2005). Data collection information can be found in Table S1.

### CryoEM data processing

GPC-I53-50 nanoparticle. The GPC-I53-50 cryoEM data was processed as described previously (Antanasijevic et al., 2020). Raw micrograph frames were aligned and dose-weighted using MotionCor2 (Zheng et al., 2017). CTF parameters were estimated with GCTF (Zhang, 2016). Initial data processing steps were performed in cryoSPARC.v2 (Punjani et al., 2017). 145,508 particles were template-picked, extracted and run through 2 rounds of 2D classification. 86,411 clean particles after 2D classification were subjected to Ab initio refinement in cryoSPARC to generate the starting model for 3D refinement steps (icosahedral symmetry was imposed). All further processing steps were performed in Relion/3.0 (Zivanov et al., 2018). 3D refinement with imposed icosahedral symmetry was run using the particles from the previous step. The refined particles were used for CTF refinement in relion (to improve the estimates for defocus and beam-tilt). To reconstruct the I53-50NP, particles from the previous step were 3D refined with a soft solvent mask around the nanoparticle core, masking out the density corresponding to the GPC trimers (icosahedral symmetry restraints imposed, local angular searches only). The resulting map was subsequently post-processed using the same soft mask to determine the B-factor (−202.1 Å^2^) and the final resolution (3.67 Å). GPC trimers were connected to I53-50A components with a flexible linker, preventing a joint analysis with the I53-50NP core. We applied localized reconstruction v1.2.0 (Ilca et al., 2015) to extract GPC trimer subparticles connected to I53-50A. 20 trimer subparticles were extracted from each GPC-I53-50NP using the pre-defined vector settings for icosahedral symmetry (--vector 0.382,0,1, --length 180). The final number of extracted GPC subparticles was 1,728,220 (20 × 86,411). The subparticle subset was subjected to two rounds of 2D classification and three rounds of 3D classification to eliminate the low-resolution classes of subparticles as well as subparticles that had issues aligning due to the signal from the nanoparticle. The final GPC subset consisted of 124,891 subparticles and was subjected to 3D auto-refinement with C3 symmetry. A soft solvent mask around the GPC trimers was applied for the refinement and post-processing steps. Final map resolution was 3.97 Å and the estimated B-factor was −123.2 Å^2^. The workflow and relevant data processing parameters are displayed in Figure S2.

GPC-I53-50A + LAVA01 Fab. The frame alignment, dose weighting and CTF estimation steps were performed as described above for the nanoparticle dataset. 416,326 particles were templated picked from the micrographs in cryoSPARC.v2 (Punjani et al., 2017) and subjected to 2D classification to remove bad particles and monomers; LAVA01 Fab has been found to induce partial trimer disassembly. 93,965 particles were selected after the 2D step and subjected to 3D refinement step in Relion/3.0. A low-pass filtered map of GPC-I53-50A trimer without the LAVA01 Fab was used as a starting model for the initial 3D refinement. The resulting particles were then subjected to another round of 3D refinement with a soft solvent mask around the GPC + LAVA01 complex. This is done to mask out the signal from the flexibly linked I53-50A nanoparticle component. This solvent mask was used for all subsequent 3D refinement and classification steps. Additionally, the particle alignment was restricted to local angular searches only (--healpix_order 3, --auto_local_healpix_order 3) to complement the usage of the solvent mask and prevent major changes in particle orientation. C1 symmetry was used for all initial 3D steps.

The dataset suffered from preferred orientation problems and it was dominated by top and bottom views of the complex. We applied an in house made program to gradually remove excess particles from over-populated views (based on Euler angle values). This was followed by iterative rounds of 3D classification (--skip_align, T=16, --sym=C1) and 3D refinement (--healpix_order 3, -- auto_local_healpix_order 3, --sym=C1). The final subset consisting of 8,480 particles was 3D refined with C3 symmetry and post-processed using the GPC + LAVA01 solvent mask. Final map resolution of the GPC + LAVA01 complex was 4.41 Å and the estimated B-factor was −71.7 Å^2^. The workflow and relevant data processing parameters are displayed in Figure S6.

### Model building and refinement

The post-processed cryoEM maps corresponding to GPC trimer antigen, I53-50NP core and GPC + LAVA01 Fab complex were used for model building and refinement. As initial models we used the published I53-50NP core (PDB ID: 6P6F; Brouwer et al., 2019) and GPC crystal structure (PDB ID: 5VK2; Hastie et al., 2017). Initial model for the LAVA01 Fab was generated using ABodyBuilder server (Leem et al., 2016). Starting models were docked into the corresponding densities and relaxed using iterative rounds of manual refinement in Coot (Casañal et al., 2020) and automated refinement in Rosetta (Wang et al., 2016). Appropriate symmetry restraints (C3 for GPC and GPC + LAVA01 Fab complex and I for I53-50NP) were applied for automated refinement steps. EMRinger (Barad et al., 2015) and MolProbity (Williams et al., 2018) were used to evaluate the refined models and generate the statistics reported in Table S2. Due to the relatively low resolution of the GPC + LAVA01 Fab complex map the amino-acid side-chains of the LAVA01 Fab were not built past Cβ.

### Rabbit immunizations

Female and naive New Zealand White rabbits (2.5–3 kg), were arbitrarily distributed in two groups of 6 rabbits and received an intramuscular immunization in each quadricep at weeks 0, 4, 16, and 28. Rabbits were immunized with either 30 µg of GPC-I53-50A or the equimolar amount presented on I53-50NPs (36 µg), both formulated in Squalene Emulsion adjuvant (Polymun, Klosterneuburg, Austria). Rabbits were sourced and housed at Covance Research Products Inc. (Denver, PA, USA) and immunizations were performed under compliance of all relevant ethical regulations for animal testing and research. The study received ethical approval from the Covance Institutional Animal Care and Use Committee with approval number C0096-019. Calculations of the dose were based on the peptidic molecular weight of the proteins which were obtained as described earlier. Bleeds were performed at weeks 0, 4, 6, 18 and 30. A larger blood draw at week 29 was also taken for isolation of peripheral blood mononuclear cells.

### Guinea pig immunizations and challenge

Naive female Hartley guinea pigs (6 weeks old) were arbitrarily distributed in two groups of 5, after which they received either 30 µg of GPC-I53-50NP adjuvanted in Squalene Emulsion adjuvant (Polymun, Klosterneuburg, Austria) or Squalene Emulsion adjuvant alone (control animals), as an intramuscular injection distributed over both quadriceps at weeks 0, 4 and 16. On day 140 (week 20) guinea pigs were challenged intraperitoneally with a targeted dose of 1 x 10^4^ PFU of guinea pig-adapted LASV Josiah strain (Safronetz et al., 2015). Animals were then monitored at least daily for fever (via transponder chip), weight loss and clinical signs of disease. Animals were euthanized via CO2 asphyxiation when humane euthanasia criteria were met, or at day 28 post-challenge. Humane euthanasia criteria were met when animals lost more than 20% of their starting weight or displayed clinical signs indicating that they had entered a moribund state (Table S5). The guinea pig studies were carried out in accordance with the recommendations per the Guide for the Care and Use of Laboratory Animals of the National Research Council. UTMB is an AAALAC-accredited institution and all animal work was approved by the Institutional Animal Care and Use Committee of UTMB under approval number 1911089. All efforts were made to minimize animal suffering and all procedures involving potential pain were performed under general anesthesia.

### Focus-forming assay

Guinea pig serum was collected in serum separator tubes, centrifuged, and frozen. On the day of the assay, a ten fold dilution series was prepared and titrated on Vero (CCL-81) cells (ATCC) with a minimal essential medium overlay containing 0.5% carboxymethylcellulose, 2% FBS, and 0.1% gentamicin in 48-well plates. Three days post-infection, plates were fixed in 10% neutral buffered formalin, and immunostained. Immunostaining was performed using anti-LASV mouse hyperimmune ascites fluid (a gift from Dr. Tom Ksiazek), and a goat anti-mouse secondary antibody (Seracare Cat# 5450-0011). Assays were developed using AEC chromagen (Enquire Bioreagents Cat# AB64252).

### Serum antibody ELISA

Half-well 96-well plates were coated overnight with Galanthus nivalis lectin (Vector laboratories) at 20 µg/mL in 0.1 M NaHCO3 pH 8.6. The next day the plates were washed three times with TBS and blocked for 30 min with Casein blocking buffer (Thermo Fisher Scientific). After washing the plates with TBS, 2 µg/mL of GPC-I53-dn5B diluted in Casein was added for 2 h. After a wash-step with TBS, three-fold dilutions of rabbit serum diluted 1:200 in TBS, 2% skimmed milk, 20% sheep serum were added to the plates. After a 2 h incubation, the plates were washed with TBS and a 1:3000 dilution of HRP-labeled goat anti-rabbit IgG (Jackson Immunoresearch) in Casein was added for 1 h. The plates were then washed five times with TBS, 0.05% Tween-20 and developing solution (1% 3,3′,5,5′-tetranethylbenzidine (Sigma-Aldrich), 0.01% H_2_O_2_, 100 mM sodium acetate and 100 mM citric acid) was added. After 1.5 min the colorimetric detection reaction was stopped by adding 0.8 M H_2_SO_4_. All procedures were performed at RT. The midpoint binding titer (ED_50_) was determined by calculating the dilution of serum that gave 50% of the maximal response from the sigmoidal binding curve. The same protocol was used for the guinea pig sera although a 1:100 start dilution was used, a 1:3000 dilution of HRP-labeled goat anti-guinea pig IgG (Thermo Fisher Scientific) and a colorimetric detection reaction of 2 min.

### B cell sorting

To generate probes for B cell sorting, biotinylated GPC-I53-50A from lineage II and IV as well as a biotinylated HCV E1E2-I53-50A (Sliepen et al., manuscript in preparation) were conjugated to the streptavidin-bound fluorophores AF647, BV421, and BB515, respectively. Conjugation was performed by incubating the biotinylated proteins for a minimum of 1 h at 4°C with the streptavidin-conjugates at a 1:2 protein to fluorochrome ratio. To saturate unconjugated streptavidin-fluorochrome complexes the fluorescent probes were next incubated for at least 10 min with 10mM biotin (GeneCopoeia) to saturate the unconjugated streptavidin-fluorochrome complexes. Week 29 rabbit PBMCs were then counted and 5×106 cells were stained for 30 min at 4 with the fluorescent probes, a viability marker (LiveDead-eF780, eBiosciences), and a rabbit PE-conjugated anti-IgG marker (Biolegend). Prior to their acquisition on the FACS-ARIA-SORP 4 laser (BD-Biosciences), cells were washed twice with FACS buffer. Viable IgG+ B cells that were negative for HCV E1E2-I53-50A and showed dual staining for both GPC-I53-50A from lineage II and IV were sorted. Analysis was performed on FlowJo v.10.7.1.

### Antibody cloning

First, a reverse transcription-PCR was performed to convert the mRNA of the lysed GPC-specific single B cells into cDNA. To do so, 6 µl of a RT-mix (200 ng) (Thermo Fisher Scientific), dNTP mix (2 mM each) (New England Biolabs), 50U Superscript III RTase, and MQ) was added to the lysed cells followed by a single cycle of 10 min at 42°C, 10 min at 25°C, 60 min at 50°C, 5 min at 95°C, and infinity 4°C. Next, the V(D)J variable regions of the cDNA were amplified by a series of PCR reactions which are distinct for heavy and light chains. The PCR mix consisted of MQ, 1x PCR reaction buffer, dNTPs (10 mM), Hotstar Plus polymerase (0.25 U; Qiagen), forward primer (25 mM), reverse primer (25 mM) (van Haaren et al., 2021; McCoy et al., 2016). For the first PCR reaction (PCR1), 13 µl of PCR mix was added to 2 µl of RT-PCR product and subjected to 5 min at 95°C, 50 cycles of [30 s at 94°C, 30 s at 58°C for light chain/48°C for heavy chain, and 1 min at 72°C], 10 min at 72°C. Next for PCR2, 13 µl of PCR mix was added to 2 µl of PCR1 product and subjected to a reaction of 5 min at 95°C, 50 cycles of [30 s at 94°C, 30 s at 55°C, and 1 min at 72°C], and 10 min at 72°C. Finally, 1 µl of PCR2 product was mixed with MQ, 1x Phusion PCR buffer, dNTPs (10 mM), forward primers (25 mM), reverse primers (25 mM), Phusion high fidelity polymerase (0.2 U; New England Biolabs) and subjected to a PCR reaction of 30 s at 98°C, 35 cycles of [5 s at 98°C, 15 s at 68°C, 20 s at 72°C], and 5 min at 72°C.

Gibson cloning was then used to integrate the amplified heavy and light chain V(D) J variable regions in mammalian cell expression vectors containing the rabbit constant regions (McCoy et al., 2016). This was done by mixing 1 µl of expression vector, 1 µl of PCR3 product, and 2 µl of Gibson mix (T5 exonuclease (0.2U; Epibio), Phusion polymerase (12.5U; New England Biolabs), Taq DNA ligase (2000U; New England Biolabs), Gibson reaction buffer (0.5 grams PEG-8000; Sigma Life Sciences), 1 M Tris/ HCl pH 7.5, 1 M MgCl2, 1 M DTT, 100 mM dNTPs, 50 mM NAD (New England Biolabs), MQ) and incubating the mix for 60 min at 50°C.

### Fab preparation

To generate LAVA01 Fab fragments, LAVA01 was subjected to a 5 h incubation at 37℃ in PBS, 10 mM EDTA, 20 mM cysteine, pH 7.4 the presence of 50 µl settled papain resin/mg of LAVA01 with papain resin. Next, Fc and non-digested mAbs were removed from the flow-through by a 2 h incubation at RT with 200 µl of protein A resin per mg of initial mAb (Thermo Scientific). Finally, the flow-through containing Fab fragments was buffer exchanged to TBS using Vivaspin filters with a 10 kDa molecular weight cutoff (GE Healthcare).

### Monoclonal antibody ELISA

A 2 µg/mL concentration of GPC-I53-dn5B diluted in Casein blocking buffer (Thermo Fisher Scientific) was added for 2 h on streptavidin-coated 96-well plates. Four-fold serial dilutions of mAbs diluted to 2.5 µg/mL in Casein were then added for 2 h. A 1:3000 dilution of HRP-labeled goat anti-rabbit IgG (Jackson Immunoresearch) in Casein was added for 1 h. Up to now, between each step, plates were washed three times with TBS. Next, plates were washed five times with TBS, 0.05% Tween-20. Colorimetric detection was performed as described above in Serum antibody ELISA. All procedures were performed at RT. The midpoint binding concentration (IC_50_) was determined by calculating the concentration of mAb that gave 50% of the maximal response from the sigmoidal binding curve.

### Generation of LASV pseudovirus

LASV pseudoviruses were generated as described previously (Robinson et al., 2016). HEK 293T cells maintained in Dulbecco’s Modified Eagle’s Medium (DMEM; Gibco) supplemented with 10% fetal calf serum (FCS), penicillin (100 U/mL), and streptomycin (100 µg/mL) were plated in 6-well plates and grown overnight to 80% confluence. The next day for each well a 1:25 dilution of Fugene in OptiMEM (final volume 250 µl; Gibco) was mixed with 0.6 µg of a full-length GPC expression plasmid and 2.4 µg of SG3Δenv (a plasmid encoding the envelope-deficient core of HIV-1) diluted in OptiMEM (final volume 250 µl; Gibcol). After a 20 min incubation at RT, the mix was added to the well. After 72 h the supernatant was harvested, sterile filtered with a 0.2 µm filter, aliquoted, and stored at −80°C. To determine virus titers, a TCID_50_ experiment was performed. TZM-bl cells were maintained in DMEM, supplemented with 10% fetal calf serum, penicillin (100 U/mL), and streptomycin (100 µg/mL) and grown overnight in a 96-well plate to a confluency of 70-80%. The next day, pseudovirus stocks were serially diluted in triplicate, incubated at RT for 1 h, and added to the TZM-bl cells. Shortly prior to addition of pseudovirus dilutions to the cells, the medium was supplemented with DEAE-dextran and Saquinavir (Sigma-Aldrich), to a final concentration of 40 µg/mL and 400 nM, respectively. Cells were then incubated at 37°C for 72 h after which they were lysed for 20 min on a shaker platform at RT by addition of Reporter Lysis Buffer (Promega). Luciferase signal was determined from cell lysates by adding the Bright-Glo Luciferase buffer (Promega) and subsequent analysis using a Glomax plate reader. The pseudovirus input for neutralization assays was determined as the dilution that gave luciferase counts of 500,000 (i.e. >10x above background).

### Pseudovirus neutralization assay

TZM-bl cells maintained in DMEM (Gibco), supplemented with 10% fetal calf serum, penicillin (100 U/mL), and streptomycin (100 µg/mL) were grown overnight in a 96-well plate to a confluency of 70-80%. The next day serial dilutions of rabbit serum or mAbs were incubated for 1hr with pseudovirus. All dilutions were performed in DMEM (Gibco), supplemented with 10% fetal calf serum, penicillin (100 U/mL), and streptomycin (100 µg/mL). The starting dilution of rabbit serum was 1:20, which was serially diluted three-fold. As guinea pig serum gave high background neutralization, IgG had to be purified from the serum as described previously (Bianchi et al., 2018). The purified IgG, diluted to the same volume as the initial serum in PBS, was diluted three-fold from a 1:20 starting dilution. For mAbs the starting concentration was 20 µg/mL or 100 µg/mL and serial dilutions were five-fold. The virus:serum/IgG/mAb mix was added to TZM-bl cells which were supplemented prior with DEAE-dextran and Saquinavir (Sigma-Aldrich), as described above. After a 72 h incubation, plates were lysed and luciferase was measured as described above. Luciferase counts were normalized to those obtained for cells infected with pseudovirus without the presence of serum/IgG/mAbs. ID_50_ and IC_50_ values were determined as the dilution/concentration where 50% inhibition of infectivity was achieved.

### Neutralization assay using authentic LASV

Neutralization assays with authentic LASV (strain Josiah, lineage IV) were performed in the BSL-4 laboratory of the Institute of Virology, Philipps University Marburg, Germany. Rabbit sera were complement inactivated for 30 min at 56 °C and diluted in a two-fold dilution series with a starting dilution of 16. Diluted sera, LAVA01, 19.7E or 37.7H (initial concentration of mAbs was 100 µg/ml), were mixed with 100 TCID_50_ virus and incubated for 60 min at 37 °C. Following incubation, Vero E6 cell suspension was added and plates were then incubated at 37°C with 5% CO_2_. Cytopathic effects (CPE) were evaluated at seven days post infection and neutralization titers were calculated as geometric mean titer (GMT) of four replicates.

### Statistical analysis

The two rabbit and guinea pigs groups were compared using a two-tailed Mann–Whitney *U*-test. Correlations were analyzed by calculating Spearman’s rank correlation coefficients. Survival comparisons were performed using a log-rank test. All statistical analyses as well as calculations of ED_50_, ID_50_, and IC_50_ values were performed using Graphpad Prism 7.0.

