## Supplemental Information for "Lassa virus glycoprotein nanoparticles elicit a neutralizing antibody that defines a new site of vulnerability"

**Table S1. CryoEM data collection information, related to figure 3 and 5**

|  | GPC-I53-50 nanoparticle | GPC-I53-50A + LAVA01 Fab |
| --- | --- | --- |
| Microscope | Titan Krios | Titan Krios |
| Voltage (kV) | 300 | 300 |
| Detector | Gatan K2 Summit | Gatan K2 Summit |
| Recording mode | Counting | Counting |
| Magnification | 29,000 | 29,000 |
| Movie micrograph pixel size | 1.03 | 1.03 |
| Dose rate (e^−^/Å^2^/s) | 4.8 | 5.3 |
| No. of frames per movie micrograph | 42 | 38 |
| Frame exposure time (ms) | 250 | 250 |
| Movie micrograph exposure time (s) | 10.5 | 9.5 |
| Total dose (e^−^/Å^2^) | 50.4 | 50.3 |
| Grid Type | QuantiFoil R 2/1 | UltrAuFoil R 1.2/1.3 |
| Under focus range (µm) | 0.6  - 1.6 | 0.6 - 1.6 |
| Number of movie micrographs | 2,337 | 3,098 |

**Table S2. Model building and refinement information, related to figure 3 and 5**

|  | GPC | I53-50 nanoparticle  (full assembly) | GPC-I53-50A + LAVA01 Fab |
| --- | --- | --- | --- |
| Final map resolution (Å) | 3.97 | 3.67 | 4.41 |
| EMDB ID | EMD-25107 | EMD-25108 | EMD-25109 |
| Residues | 1089 | 21060 | 1728 |
| Amino-acids | 1014 | 21060 | 1641 |
| Carbohydrates | 75 | 0 | 87 |
| RMSD Bond Length (4𝛔) | 0.023 | 0.020 | 0.021 |
| RMSD Bond Angles (4𝛔) | 1.733 | 1.688 | 1.971 |
| Ramachandran |  |  |  |
| Outliers (%) | 0.0 | 0.0 | 0.0 |
| Allowed (%) | 1.8 | 2.3 | 1.9 |
| Favored (%) | 98.2 | 97.7 | 98.1 |
| Rotamer Outliers (%) | 0.0 | 0.0 | 0.0 |
| Clash Score | 1.47 | 0.19 | 1.10 |
| MolProbity Score | 0.88 | 0.64 | 0.82 |
| EMRinger Score | 2.57 | 2.81 | 1.37 |
| PDB ID | 7SGD | 7SGE | 7SGF |

**Tabel S3. Midpoint pseudovirus neutralization titers from rabbits that received GPC-I53-50A or GPC-I53-50NPs, tested against a panel of LASV-pseudoviruses, related to figure 5**

ID_50_ values, i.e. the serum dilution at which infectivity was inhibited by 50%, are shown and color coded: white = no neutralization, ID_50_ < 20; yellow = weak neutralization, 20 > ID_50_ >100; orange = moderate neutralization, 101 > ID_50_ > 1000; red = strong neutralization, ID_50_ > 1000. BG505 = HIV pseudovirus (negative control).

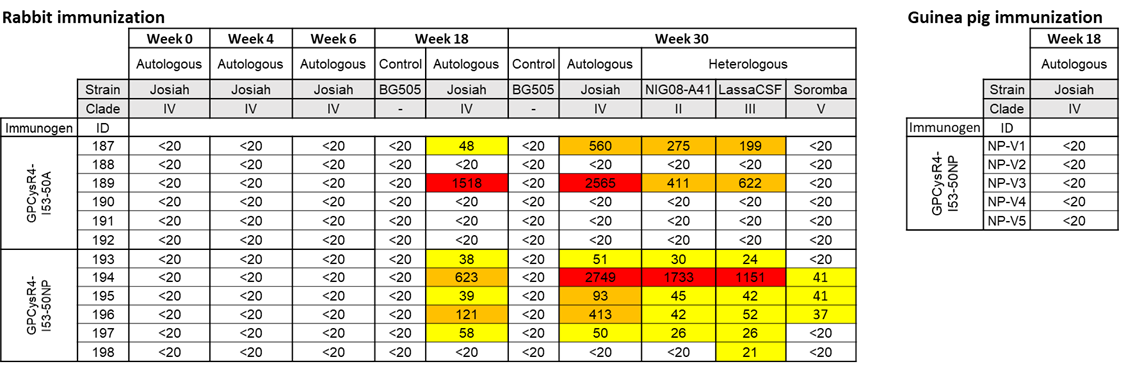

**Tabel S4. Endpoint neutralization titers from rabbits that received GPC-I53-50A or GPC-I53-50NPs, tested against authentic LASV, related to figure 5**

The virus neutralization titer is calculated as the geometric mean titers (GMT) of the reciprocal value of the last serum dilution at which inhibition of the cytopathic effect on infected Vero E6 cells is detectable. The initial dilution was 1:16 so a titer of 8 was noted when no inhibition was observed.

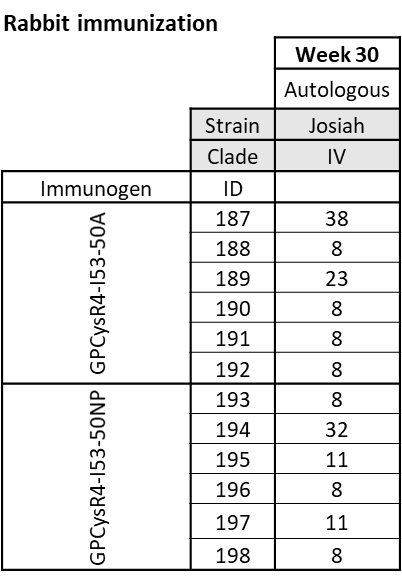

**Tabel S5. Midpoint pseudovirus neutralization titers from guinea pigs that received GPC-I53-50NPs, tested against autologous LASV-pseudovirus, related to figure 5**

ID_50_ values, i.e. the purified IgG dilution at which infectivity was inhibited by 50%, are shown and color coded: white = no neutralization, ID_50_ < 20.

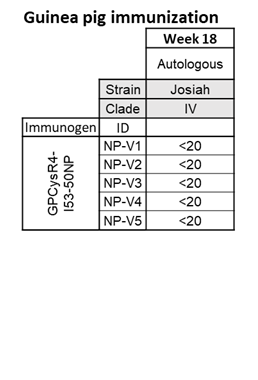

**Table S6. Scoring scheme to assess the health of guinea pigs post-challenge, related to figure 5**

| **Score (1-4)** | **Description of Animal** |
| --- | --- |
| **1** | Healthy |
| **2** | Ruffled fur and Hunched posture (Triggers 2^nd^ observation) |
| **3** | A score of 2 plus 1 additional clinical sign such as, Lethargy, Orbital tightening, and/or >15% weight loss (Triggers 3rd observation) |
| **4** | A score of 3 plus 1 additional clinical sign such as, Refusal to stampede, or any neurologic signs (rectal prolapse, seizures, tremors, head tilt, paralysis, etc.) OR >20% weight loss - Immediate Euthanasia |

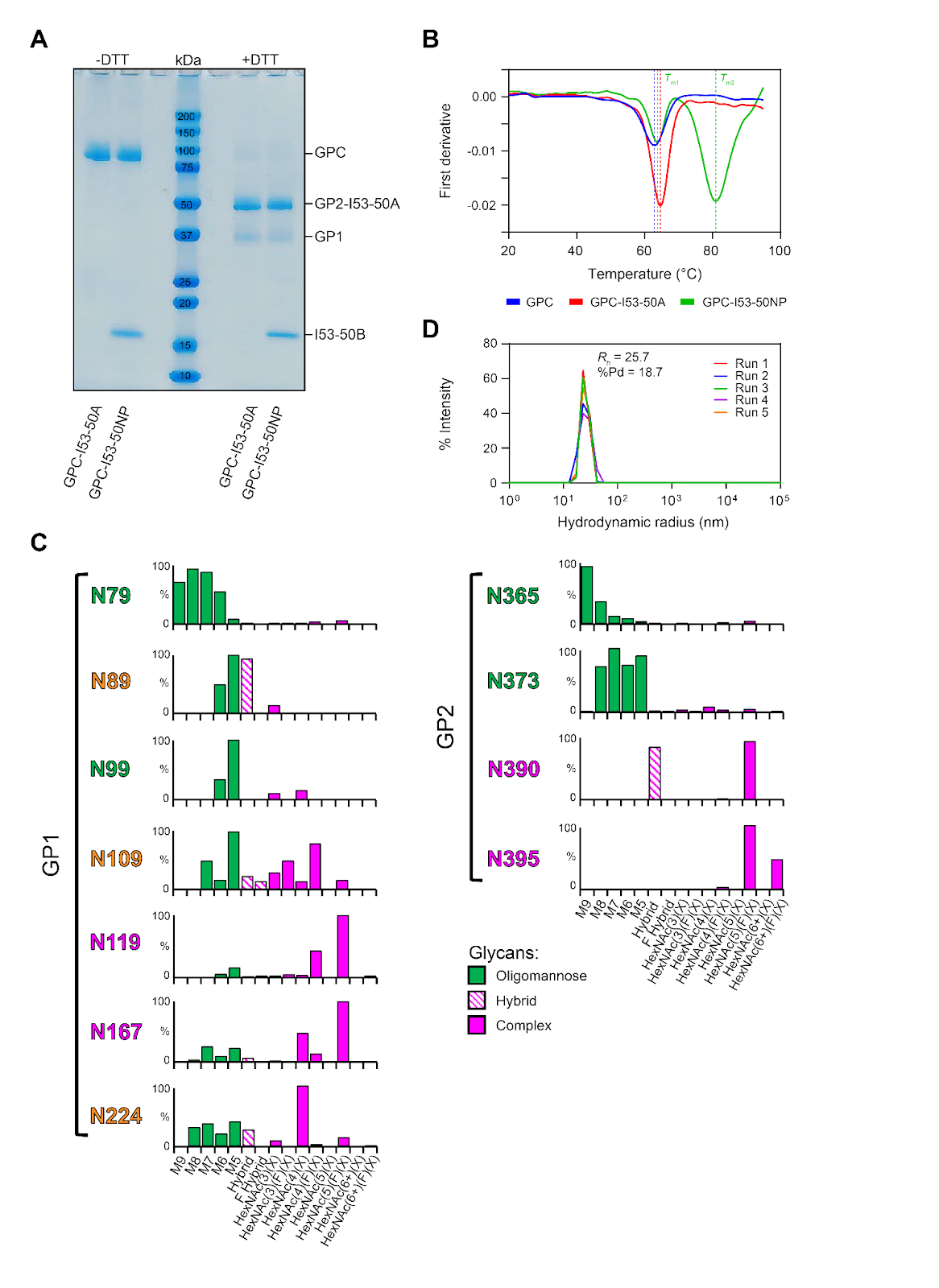

**Fig. S1. Biophysical characterization of GPC-I53-50A and GPC-I53-50NPs, related to figure 1 and 2**

(A) SDS-PAGE analysis of GPC-I53-50A and GPC-I53-50NP in the absence (-DTT) and presence (+DTT) of dithiothreitol (DTT). (B) NanoDSF curves of GPC, GPC-I53-50A, and GPC-I53-50NP.  The dotted lines indicate the melting tempera-tures (*T*_m_), which is defined as the temperature where 50% of the protein is unfolded. GPC-I53-50NP has two melting temperatures; one for unfolding of GPC (*T*_m1_) and one for the I53-50NP core (*T*_m2_). Representative melting curve of at least two technical replicates is shown. (C) Site-specific glycan distribution of N-linked glycans on GPC-I53-50A as determined by LC-MS. The bar graphs represent the relative quantities of each glycan group with oligomannose-type glycan series (M9 to M5; Man9GlcNAc2 to Man5GlcNAc2) (green), a fucosylated and fucosylated hybrid glycans (Hybrid & F Hybrid) (dashed pink) and complex glycans grouped according to fucosylation and the number of antennae (HexNAc(3)(X) to HexNAc(6+)(F)(X)) (pink). (D)  DLS data of GPC-I53-50NPs with the average polydispersity (%Pd) and hydrodynamic radius (*R*_h_) of 5 runs shown. An overlay of each run is shown. A %Pd < 15 is considered a monodisperse population.

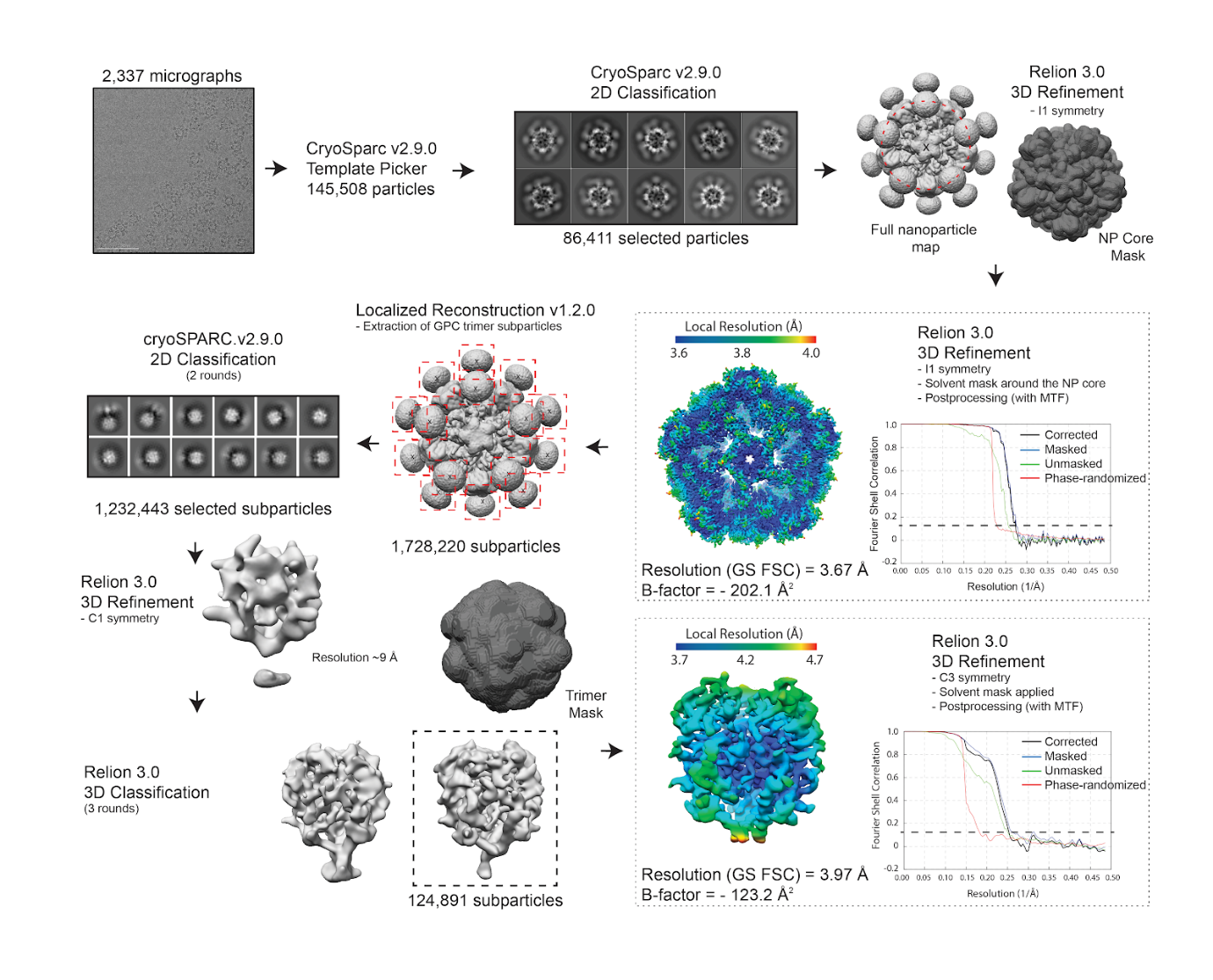
**Fig. S2. CryoEM data processing workflow for GPC-I53-50 nanoparticles, related to figure 3**

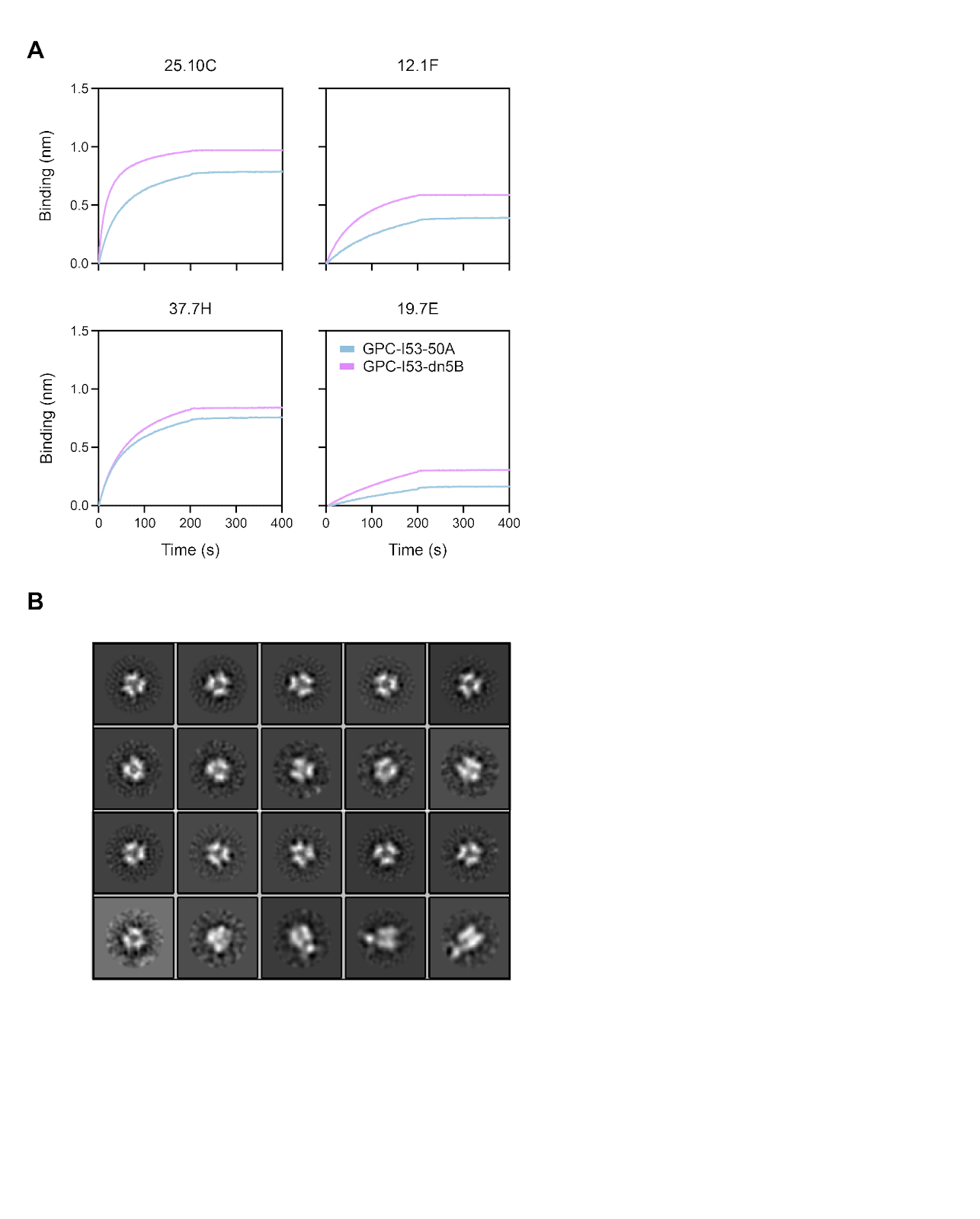

**Fig. S3. Antigenic and structural characterization of GPC-dn5B, related to figure 4**

(A) Sensorgrams from BLI experiments comparing the binding of GPC-specific mAbs 25.10C, 12.1F, 37.7H, and 19.7E by GPC-I53-50A and GPC-dn5B. (B) 2D-class averages from nsEM with GPC-dn5B.

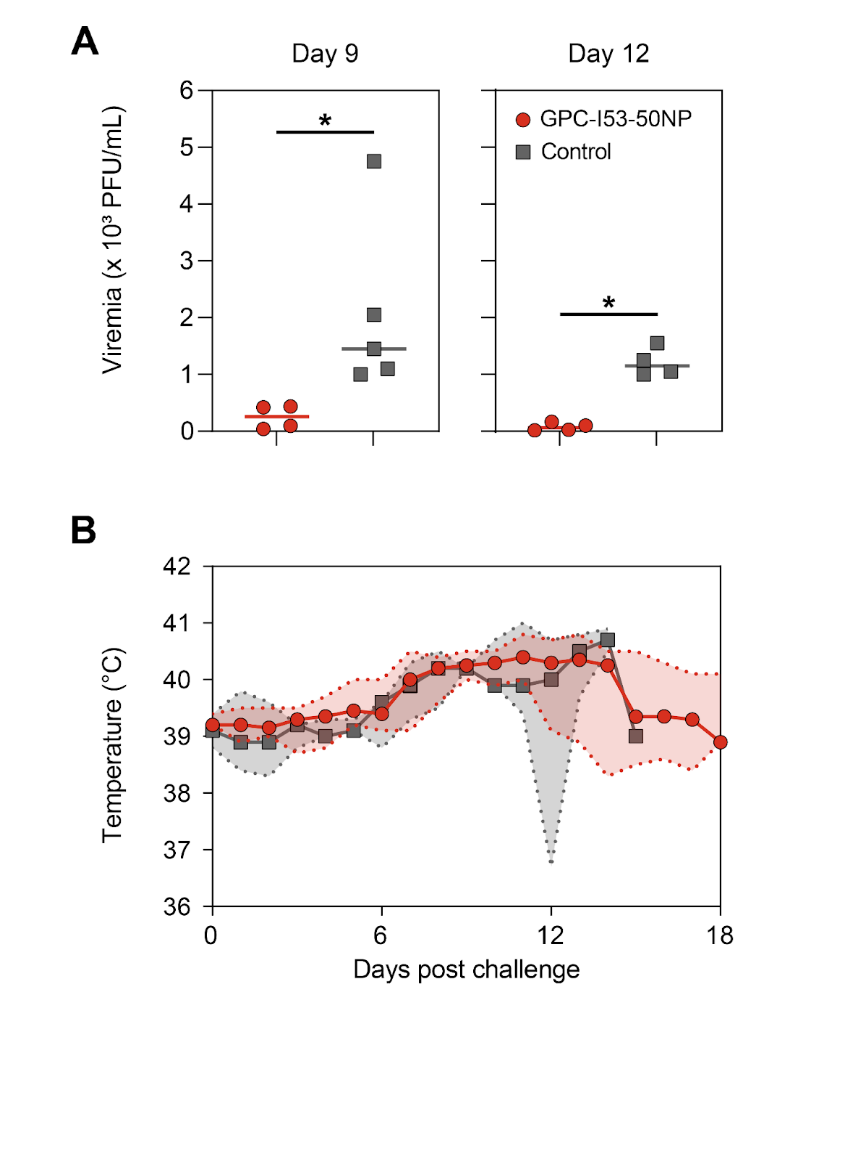

**Fig. S4. Virema, body temperature and weight alteration in vaccinated and control guinea pigs after challenge, related to figure 4**

(A) Median RNA viral loads in vaccinated and control guinea pigs after challenge at week 9 and week 12. The shaded area indicates the range. Statistical differences between two groups (week 9: *n* = 4 for vaccinated, *n* = 5 for controls; week 12: *n* = 4 for vaccinated and controls) were determined using two-tailed Mann–Whitney U-tests (**p* < 0.05). (B) Median body temperature over time of vaccinated and control guinea pigs after challenge. The shaded area indicates the range.

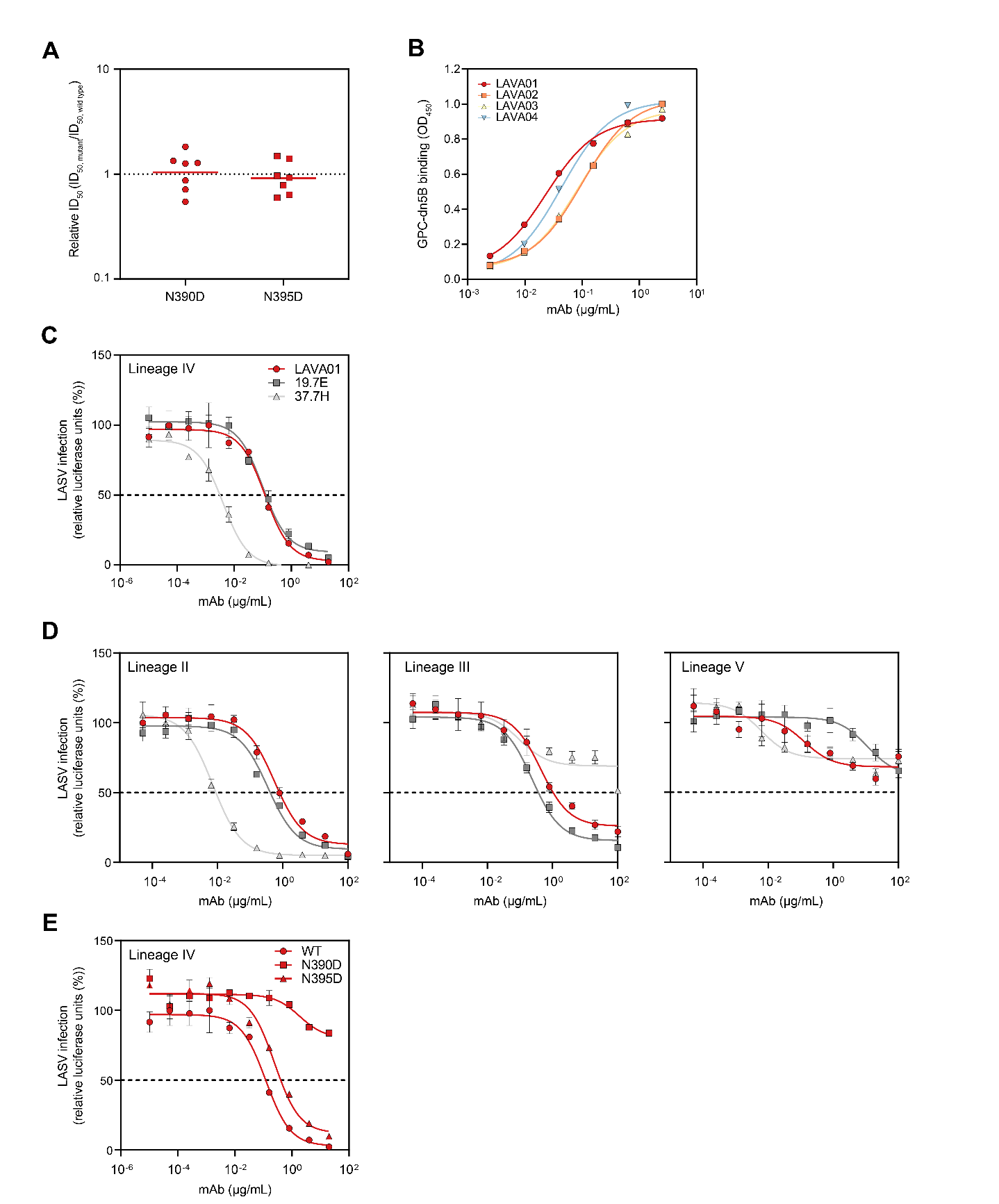

**Fig. S5. Dependency of rabbit serum and LAVA01 neutralization on N390 and N395 glycans, binding of isolated mAbs to GPC, and comparison of LASV pseudovirus neutralization between LAVA01, 19.7E and 37.7H, related to figure 5**

(A) Relative ID_50_ for the N390D and N395D pseudovirus mutant relative to the parental lineage IV (Josiah) pseudovirus for all rabbit sera that showed an autologous neutralization ID_50_ titer >20. The dotted line indicates a RID_50_ of 1, i.e. the mutation has no effect on neutralization.  (B) Binding of LAVA01-LAVA04 to GPC-dn5B as determined by ELISA. (C) Neutralization of lineage IV (Josiah) pseudovirus by LAVA01, 19.7E, and 37.7H, as indicated in the legend. (D) Neutralization of lineage II (NIG08-A41; left), III (CSF; middle), and V (Bamba; right) pseudovirus by LAVA01, 19.7E, and 37.7H. The mean and SEM of at least two technical replicates are shown. (E) Neutralization curves of N390D or N395D pseudovirus mutants compared to the parental lineage IV pseudovirus (WT) by LAVA01. (C)-(E) The dotted line indicates 50% neutralization.

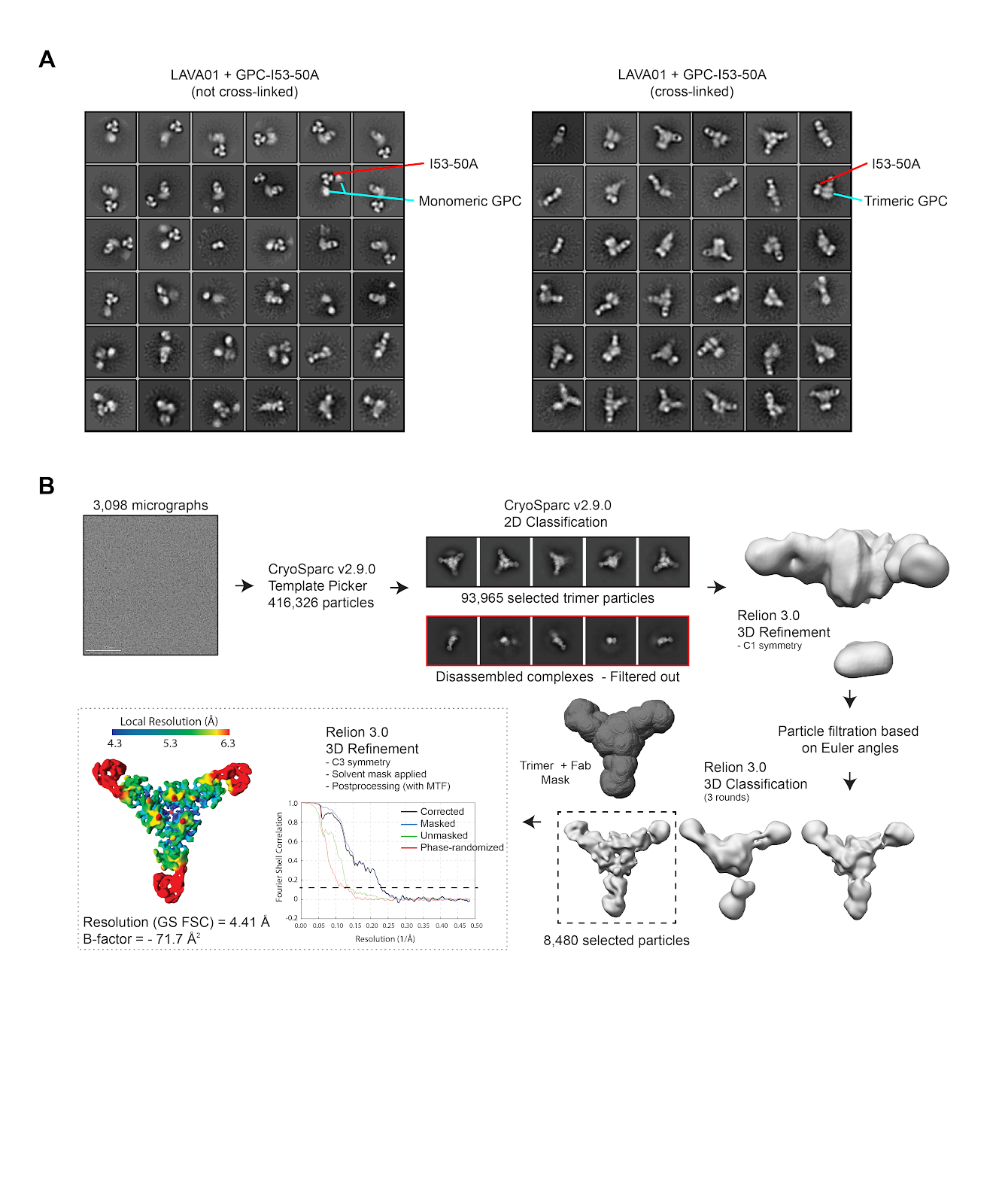
**Fig. S6. CryoEM data processing workflow for complexes of chemically cross-linked GPC-I53-50A with LAVA01, related to figure 5**

(A) 2D-class averages from nsEM with complexes of LAVA01 with GPC-I53-50A (left) and chemically cross-linked GPC-I53-50A (right). The I53-50A component and GPC are indicated. (B) CryoEM data processing workflow for complexes of LAVA01 with chemically cross-linked GPC-I53-50A.
